# Mapping the *Arabidopsis thaliana* proteome in PeptideAtlas and the nature of the unobserved (dark) proteome; strategies towards a complete proteome

**DOI:** 10.1101/2023.06.01.543322

**Authors:** Klaas J. van Wijk, Tami Leppert, Zhi Sun, Alyssa Kearly, Margaret Li, Luis Mendoza, Isabell Guzchenko, Erica Debley, Georgia Sauermann, Pratyush Routray, Sagunya Malhotra, Andrew Nelson, Qi Sun, Eric W. Deutsch

## Abstract

This study describes a new release of the *Arabidopsis thaliana* PeptideAtlas proteomics resource providing protein sequence coverage, matched mass spectrometry (MS) spectra, selected PTMs, and metadata. 70 million MS/MS spectra were matched to the Araport11 annotation, identifying ∼0.6 million unique peptides and 18267 proteins at the highest confidence level and 3396 lower confidence proteins, together representing 78.6% of the predicted proteome. Additional identified proteins not predicted in Araport11 should be considered for building the next Arabidopsis genome annotation. This release identified 5198 phosphorylated proteins, 668 ubiquitinated proteins, 3050 N-terminally acetylated proteins and 864 lysine-acetylated proteins and mapped their PTM sites. MS support was lacking for 21.4% (5896 proteins) of the predicted Araport11 proteome – the ‘dark’ proteome. This dark proteome is highly enriched for certain (*e.g.* CLE, CEP, IDA, PSY) but not other (*e.g.* THIONIN, CAP,) signaling peptides families, E3 ligases, TFs, and other proteins with unfavorable physicochemical properties. A machine learning model trained on RNA expression data and protein properties predicts the probability for proteins to be detected. The model aids in discovery of proteins with short-half life (*e.g.* SIG1,3 and ERF-VII TFs) and completing the proteome. PeptideAtlas is linked to TAIR, JBrowse, PPDB, SUBA, UniProtKB and Plant PTM Viewer.

## INTRODUCTION

*Arabidopsis thaliana* (Arabidopsis) was established as a universal plant model system in the 1980s as a means of advancing the plant science field (Meinke et al., 1998; Koornneef and Meinke, 2011). The power of Arabidopsis as an experimental model system to discover novel gene functions and molecular pathways was first demonstrated using loss-of function mutants in the photorespiratory pathway (Somerville and Ogren, 1980, 1982). Since then, the field of plant biology, and specifically plant molecular biology and genetics, has expanded enormously and produced a wealth of knowledge and understanding of plants (Parry et al., 2020; Provart et al., 2021). A well-organized Arabidopsis community with powerful public resources is facilitating and accelerating new discoveries in Plant Biology (Parry et al., 2020; Alex Mason et al., 2021).

Arabidopsis also has been established as a model for analysis of its proteome in particular because mass spectrometry (MS) based proteomics immensely benefits from having a well-annotated genome with a robust set of predicted proteins (van Wijk et al., 2021). A poorly annotated genome and poorly predicted proteins diminish the ability to carry out quantitative proteome analyses and determine the rich complexity of post-translational modifications (PTMs), including the assignment of PTMs to specific amino acid residues. A range of plant proteome databases by individual labs has been developed, mostly for Arabidopsis proteins, often focused on a particular aspect of plant proteomics, such as subcellular compartments (San Clemente and Jamet, 2015; Salvi et al., 2018), protein location (SUBA and PPDB) (Sun et al., 2009; Tanz et al., 2013), or PTMs (Schulze et al., 2015; Willems et al., 2019). A comprehensive Arabidopsis proteome database (ATHENA) was released to allow mining of a large-scale experimental proteome dataset involving multiple tissue types as published in (Mergner et al., 2020). In 2021, we launched the first release of the Arabidopsis PeptideAtlas to addressentral questions about the Arabidopsis proteome, such as experimental evidence for accumulation of proteins, their approximate relative abundance, the significance of protein splice forms, and selected PTMs, (van Wijk et al., 2021) (https://peptideatlas.org/builds/arabidopsis/). Species-specific PeptideAtlas resources have also been developed for non-plant species including human (Omenn et al., 2021), various animals such pigs (Hesselager et al., 2016), chicken (McCord et al., 2017), fish (Nissa et al., 2022), different yeast species (King et al., 2006; Gunaratne et al., 2013) and bacteria (Malmstrom et al., 2009; Michalik et al., 2017; Reales-Calderon et al., 2021). Each PeptideAtlas is based on published MSMS datasets collected through the ProteomeXchange Consortium (Deutsch et al., 2023) and reanalyzed through a uniform processing pipeline. In the case of the Arabidopsis PeptideAtlas, we are particularly keen to annotate the metadata associated with the raw MS data and to link all peptide identifications to spectral, technical and biological metadata. These metadata are critical to determine cell-type or sub-cellular specific protein accumulation patterns and help accomplish the long-term goal of the Arabidopsis community to develop a detailed Arabidopsis Plant Cell Atlas (Plant Cell Atlas et al., 2021).

The current study describes the second PeptideAtlas release, which adds an additional 63 ProteomeXchange datasets (PXDs) containing 102 million MSMS spectra to the first release in 2021. The objectives for this second release were to map peptides to proteins that were not identified in the 1st release and to extend sequence coverage of already identified proteins. In addition, the second release would provide deeper coverage for protein phosphorylation and N-terminal and lysine acetylation, and now also includes PXDs that included specific enrichment workflows for ubiquitinated proteins (Walton et al., 2016; Grubb et al., 2021). To try to increase the detection or sequence coverage of proteins, we employed four criteria for the selection of new PXDs: i) PXDs of specific cell types or specialized subcellular fractions, ii) PXDs that concern specific protein complexes or protein affinity enrichments, iii) PXDs that are enriched for specific post-translational modifications, and iv) PXDs that appear to have very high dynamic resolution and sensitivity by using the latest technologies in mass spectrometry and/or sample fractionation. The new PeptideAtlas release now maps peptides to 78.6% of the predicted Arabidopsis proteome, with each mapped peptide connected to the metadata and spectrum matches. With the ultimate goal to identify the complete Arabidopsis predicted proteome, we investigated why 21% of the predicted Arabidopsis proteome was not yet observed in this new build. A significant portion of these unobserved proteins have physicochemical properties that should impede detection by MS (*e.g*. very small, very hydrophobic). Other unobserved proteins likely accumulate under highly specific conditions or cell-types and/or have low cellular abundance. Here we used large scale RNA-seq data sets for Arabidopsis to determine mRNA expression patterns for these unobserved proteins sampling across many tissue- and cell types, developmental stages, as well as biotic and abiotic stress conditions. We developed machine learning models based on these mRNA expression features and physicochemical protein properties to calculate the probability for each protein to be detected. GO enrichment analysis showed over-representation of specific functions in the dark proteome, *e.g.* E3 ligases and signaling peptides. The machine learning model outputs will help design optimal and targeted experimental strategies to detect these unobserved proteins. Finally, this second PeptideAtlas release including its associated metadata and our machine learning output provides an ideal platform to contribute to a community Arabidopsis proteome cell atlas (Plant Cell Atlas et al., 2021; Birnbaum et al., 2022) and also contribute to the ongoing reannotation of the Arabidopsis genome (tinyurl.com/Athalianav12). The new PeptideAtlas release is integrated into TAIR (https://www.arabidopsis.org/) and linked to JBrowse (https://jbrowse.arabidopsis.org), PPDB (Sun et al., 2009), SUBA (Hooper et al., 2017), UniProtKB (UniProt, 2023) and Plant PTM Viewer (Willems, 2022).

## MATERIALS AND METHODS

### Selection and downloads of ProteomeXchange submissions

Raw files for the selected PXDs were downloaded from ProteomeXchange (http://www.proteomexchange.org/) repositories. Supplemental Table Data Set 1 provides detailed information about the 63 newly selected PXDs, as well as the 52 PXDs that were part of the first build; this includes information about instrument, sample (*e.g.* subcellular proteome, plant organ), number of raw files and MSMS spectra (searched and matched), identified proteins and peptides, submitting lab and associated publication, as well as several informative key words.

### Extraction and annotation of metadata

For each selected dataset, we obtained information associated with the submission, and the publication if available, to determine search parameters and provide meaningful tags that describe the samples in some detail. These tags are visible for the relevant proteins in the PeptideAtlas interface. If needed, we contacted the submitters for more information about the raw files. All collected metadata are stored in our annotation system as previously described (van Wijk et al., 2021). These metadata can be viewed for each identified protein in PeptideAtlas.

### Assembly of protein search space

We assembled a comprehensive protein search space comprising the predicted *Arabidopsis* protein sequences from i) Araport11 (Cheng et al., 2017), ii) TAIR10 (Lamesch et al., 2012), iii) UniProtKB (UniProt, 2020), iv) RefSeq (https://www.ncbi.nlm.nih.gov/refseq) (Li et al., 2021), v) from the repository ARA-PEPs (Hazarika et al., 2017) with 7901 small Open Reading Frames (sORFs), 16809 low molecular weight peptides and proteins (LWs; between 26 and 250 aa; median 37 aa), as well as 607 novel stress-induced peptides (SIPs) most of which are currently not annotated in TAIR10 or Araport11, vi) from Dr Eve Wurtele (Iowa State University) assembled based on RNA-seq data, vii) GFP, RFP and YFP protein sequences commonly used as reporters and affinity enrichments, viii) 116 contaminant protein sequences frequently observed in proteome samples (*e.g.* keratins, trypsin, BSA) (https://www.thegpm.org/crap/). This search space is quite similar as for the first PeptideAtlas release, except that the UniProtKB and RefSeq contributions were updated to the latest version as of 2021-04. Also added was the complete set of predicted protein sequences for the 950 Araport11 pseudogenes (1240 gene models) that we generated through 3-frame translation (the pseudogene sequences have transcription direction, but no frame).

We also included an update on the plastid- and mitochondrial-encoded proteins to address redundancies in plastid- and mitochondrial ATGC and ATMG identifiers, and inclusion of protein sequences for those plastid- and mitochondrial encoded proteins that are predicted to be affected by RNA editing. For the mitochondrial-encoded proteins, we included 420 editing sites in 29 mitochondrial-encoded proteins and two ORFs, most of which are described in (Sloan et al., 2018) whereas we included edited sequences for 17 plastid-encoded proteins that included 31 amino acid changes and generation of one start methionine. These organellar-encoded sequences included unedited sequences, completely edited sequences, and if editing sites were sufficiently close together to appear in a single peptide, we also include all permutations of edits and non-edits. This resulted in the addition of 10,368 sequences for plastid- and mitochondrial encoded variants to the search database. In a forthcoming study (van Wijk et al, in preparation), we will provide details on the annotation and redundancy of plastid- and mitochondrial encoded proteins, the expression of organellar ORFs, and the impact of RNA editing.

### The Trans-Proteomic Pipeline (TPP) data processing pipeline

For all selected datasets, the vendor-format raw files were downloaded from the hosting ProteomeXchange repository, converted to mzML files (Martens et al., 2011) using ThermoRawFileParser (Hulstaert et al., 2020) for Thermo Fisher Scientific instruments or the msconvert tool from the ProteoWizard toolkit (Chambers et al., 2012) for SCIEX wiff files, and then analyzed with the TPP (Keller et al., 2005; Deutsch et al., 2015) version 6.2.0 (Deutsch et al., 2023). The TPP analysis consisted of sequence database searching with either Comet (Eng and Deutsch, 2020) for LTQ-based fragmentation spectra or MSFragger 3.2 (Kong et al., 2017) for higher resolution fragmentation spectra and post-search validation with several additional TPP tools as follows: PeptideProphet (Keller et al., 2002) was run to assign probabilities of being correct for each peptide-spectrum match (PSM) using semi-parametric modeling of the search engine expect scores with z-score accurate mass modeling of precursor m/z deltas. These probabilities were further refined via corroboration with other PSMs, such as multiple PSMs to the same peptide sequence but different peptidoforms or charge states, using the iProphet tool (Shteynberg et al., 2011).

For datasets in which trypsin was used as the protease to cleave proteins into peptides, two parallel searches were performed, one with full tryptic specificity and one with semi-tryptic specificity. The semi-tryptic searches were carried out with the following possible variable modifications (5 max per peptide for Comet and 3 for MSFragger): oxidation of Met or Trp (+15.9949), acetylation of Lys (+42.0106), peptide N-terminal Gln to pyro-Glu (-17.0265), peptide N-terminal Glu to pyro-Glu (-18.0106), deamidation of Asn or Gln (+0.9840), peptide N-term acetylation (+42.0106), and if peptides were specifically affinity enriched for phosphopeptides, also phosphorylation of Ser, Thr or Tyr (+79.9663). For the fully tryptic searches, we also added oxidation of His (+15.9949) and formylation of peptide N-termini, Ser, or Thr (+27.9949)] - we deliberately restricted these latter PTMs to only full tryptic (rather than also allowing semi-tryptic) to reduce the search space and computational needs. Formylation is a very common chemical modification that occurs in extracted proteins/peptides during sample processing, whereas His oxidation is observed less frequently, but nevertheless at significant levels (Verrastro et al., 2015; Hawkins and Davies, 2019). In both semi-tryptic and full tryptic searches, fixed modifications for carbamidomethylation of Cys (+57.0215) if treated with reductant and iodoacetamide (or other alkylating reagents) and isobaric tag modifications (TMT, iTRAQ) were applied as appropriate. Both variable and fixed modifications were applied to dimethyl labeled datasets as appropriate. Four missed cleavages were allowed (RP or KP do not count as a missed cleavage). Several PXDs were generated using other proteases (GluC, ArgC, Chymotrypsin); these data sets were processed similarly to those generated by trypsin with the exception that the relevant enzyme was chosen. Some of the datasets derived from analysis of extracted peptidomes in which ‘no protease treatment’ was used and these datasets were searched with ‘no enzyme’.

### PeptideAtlas Assembly

In order to create the combined PeptideAtlas build of all experiments, all datasets were thresholded at an iProphet probability that yields the model-based PSM FDR of 0.0008. The exact probability varied from experiment to experiment depending on how well the modeling can separate correct from incorrect. This probability threshold was typically greater than 0.99. As more and more experiments are combined, the total FDR increases unless the threshold is made more stringent (Deutsch et al., 2016). Throughout the procedure, decoy identifications were retained and then used to compute final decoy-based FDRs. The decoy count-based PSM-level FDR was 0.0001 (8001 decoy PSMs out of 70 million), peptide sequence-level FDR is 0.001 (728 decoy sequences out of 596,839), and the final canonical protein-level FDR was 0.0005 (10 decoy proteins out of 18,267 with canonical status). Among proteins with lesser status (weak, insufficient evidence, etc.) there are 645 decoys out of 21,854 yielding an FDR of 0.03. Because of the tiered system, high quality MSMS spectra that were matched to a peptide are never lost, even if a single matched peptide by itself cannot confidently identify a protein.

### Protein identification confidence levels and classification

Proteins were identified at different confidence levels using standardized assignments to different confidence levels based on various attributes and relationships to other proteins. The highest level is canonical and lowest is ‘not detected’. In between are various levels of uncertain and redundant proteins; this tier system was described in detail in (van Wijk et al., 2021) and will not be repeated here.

### Handling of gene models and splice forms

The 27655 protein coding genes in Araport11 are represented by 48359 gene models (transcript isoforms), which are identified by the digit after the AT identifier (*e.g.* AT1G10000.1). We refer to the translations of these gene models as protein isoforms. Most protein isoforms are either identical or very similar (differing only a few amino acid residues often at the N-or C-terminus). It is often hard to distinguish between different protein isoforms due to the incomplete sequence coverage inherent to most MS proteomics workflows. For the assignment of canonical proteins (at least two uniquely mapping peptides identified), we selected by default only one of the protein isoforms as the canonical protein; this was the ‘.1’ isoform unless one of the other isoforms had a higher number of matched peptides. However, if other protein isoforms did have detected peptides that are unique from the canonical protein isoform (*e.g.* perhaps due to a different exon), then they can be given canonical (tier 1) or less confident tier status depending on the nature of the additional uniquely mapping peptides (length and numbers). If the other protein isoforms do not have any uniquely mapping peptides amongst all protein isoforms (for that gene), then they are classified as redundant.

### Integration of PeptideAtlas results in other web-based resources

PeptideAtlas is accessible through its web interface at https://peptideatlas.org. Furthermore, direct links are provided between PeptideAtlas and PPDB (http://ppdb.tc.cornell.edu/), UniProtKB (https://www.uniprot.org/), TAIR (https://www.arabidopsis.org/), Plant PTM Viewer (https://www.psb.ugent.be/webtools/ptm-viewer/), PhosPhAt (http://phosphat.uni-hohenheim.de/), SUBA5 (https://suba.live/), ATHENA (http://athena.proteomics.wzw.tum.de:5002/master_arabidopsisshiny/), and several more. Links to matched peptide entries in PeptideAtlas are available in the Arabidopsis annotated genome through a specific track in JBrowse at https://jbrowse.arabidopsis.org.

### Protein physicochemical properties and functions

To characterize the canonical and unobserved proteomes, physicochemical properties were calculated or predicted using various web-based tools. These include: protein length, mass, GRAVY index, isoelectric point (pI), number of transmembrane domains (http://www.cbs.dtu.dk/services/TMHMM) and sorting sequences for the ER, plastids and mitochondria (http://www.cbs.dtu.dk/services/TargetP-1.0/).

### Assembly and quality control filtering of the RNA dataset

13,673 single and paired end RNA-seq datasets from (Palos et al., 2022) were run through featureCounts (Liao et al., 2014) to count reads aligning to each of the 27,655 Arabidopsis genes. Lower quality datasets were filtered out based on a minimum total read count (5,000,000), eliminating 7,994 datasets. Transcripts Per Million (TPM) expression values were calculated for the remaining 5,679 datasets. Genes for which expression above zero TPM was not detected in any of the remaining datasets were removed, eliminating 398 genes. The median TPM value for each dataset was then calculated and used as the threshold for the identification of expressed genes within the dataset. Six datasets had a median of 0 and were removed from the analysis. Furthermore, 345 protein-coding genes were never expressed above the median. These genes and the datasets in which they are transcribed are described in the Supplemental Data Set 2.

### Machine learning - Developing Classification Models

The artificial neural network (ANN) model and the random decision forest (RDF) models are trained using Python 3.8.10 with TensorFlow 2.12.0 and TensorFlow Decision Forests 1.3.0 respectively. The input file used for both models is derived from a dataset containing 23,674 Arabidopsis canonical and unobserved proteins and their attributes. Each entry in the dataset includes the protein’s identifier, gene symbol, the chromosome on which it is found, its status of being “canonical” or “not observed”, number of recorded observations, a short description, molecular weight, gravy, pI, percentage of RNA-seq datasets detecting it, and highest TPM. Only the last five columns are selected for training in the input file. To accommodate the prediction tools, the status is denoted by a 1 or 0 that represents “canonical” or “not observed” respectively. All Python code used for the modeling and the output files are available on GitHub at https://github.com/PlantProteomes/ArabidopsisDarkProteome.

## RESULTS & DISCUSSION

### Selection of PXDs

In July 2022, there were ∼630 PXDs for Arabidopsis publicly available in ProteomeXchange (Figure 1A) most of which were submitted through PRIDE (Perez-Riverol et al., 2018; Perez-Riverol et al., 2022) (89%) and the remainder through MassIVE (Pullman et al., 2018), JPOST (Moriya et al., 2019), iProX (Ma et al., 2019) or Panorama Public (Sharma et al., 2018). For most of these PXDs (84%) the MS data were acquired using an Orbitrap type instrument (*e.g.*, Q Exactive models, LTQ-Orbitrap Velos/XL/Elite, Orbitrap Fusion Lumos) and the remainder a variety of instruments (*e.g.* TripleTOF and Maxis/Impact II) from different vendors (Figure 1B). For build 2, we selected 63 new PXDs and analyzed these together with all 52 datasets from build 1. Table 1 summarizes key information for all 115 selected PXDs in build 2; additional information can be found in Supplemental Data Set 1. These new PXDs were selected because they appeared the most promising to identify new proteins and selected PTMs, as well as increase sequence coverage of proteins already identified at lower (non-canonical) confidence levels. For example, the selected PXDs concerned specific protein complexes (*e.g*. mitochondrial ribosomes PXD010324 (Waltz et al., 2019)), proximity labeling to target subcellular complexes (*e.g*. the nuclear pore complex PXD015919 (Huang et al., 2020), and subcellular localizations (*e.g*. clathrin-coated vesicles PXD026180 (Dahhan et al., 2022) that were underrepresented. We also selected two large studies involving affinity-enrichment for ubiquitination (Walton et al., 2016; Grubb et al., 2021), a study enriching for SUMOylated proteins (Rytz et al., 2018), as well as additional PXDs involving n-terminal or lysine acetylation or phosphorylation. We do note that most PXDs involved the Col-0 ecotype (for which most community resources are available), but one study used ecotype *Wassilewskija* (Ws) and six studies used cell cultures generated from *Landsberg erecta* (Ler). Because of the complexities of data processing and control of the overall false discovery rate (FDR), we excluded data sets obtained through data independent acquisition (DIA), targeted MS (MRM or SRM) and only considered data dependent acquisition (DDA). However, we did include stable isotope labeled (multiplexed) proteome datasets, including isobaric iTRAQ and TMT (Rauniyar and Yates, 2014; Chen et al., 2021), dimethyl labeling (Hsu et al., 2003), as well as N-terminomics datasets using TAILS (Kleifeld et al., 2011) or COFRADIC (Gevaert et al., 2003). Finally, we also considered mass spectrometer type with preference for Orbitrap-type instruments (Thermo) because of their high mass accuracy, ease of reprocessing, and because ∼84% of all available PXDs in ProteomeXchange used such Orbitrap instruments (Table 1; Figure 1B).

**Figure 1.**
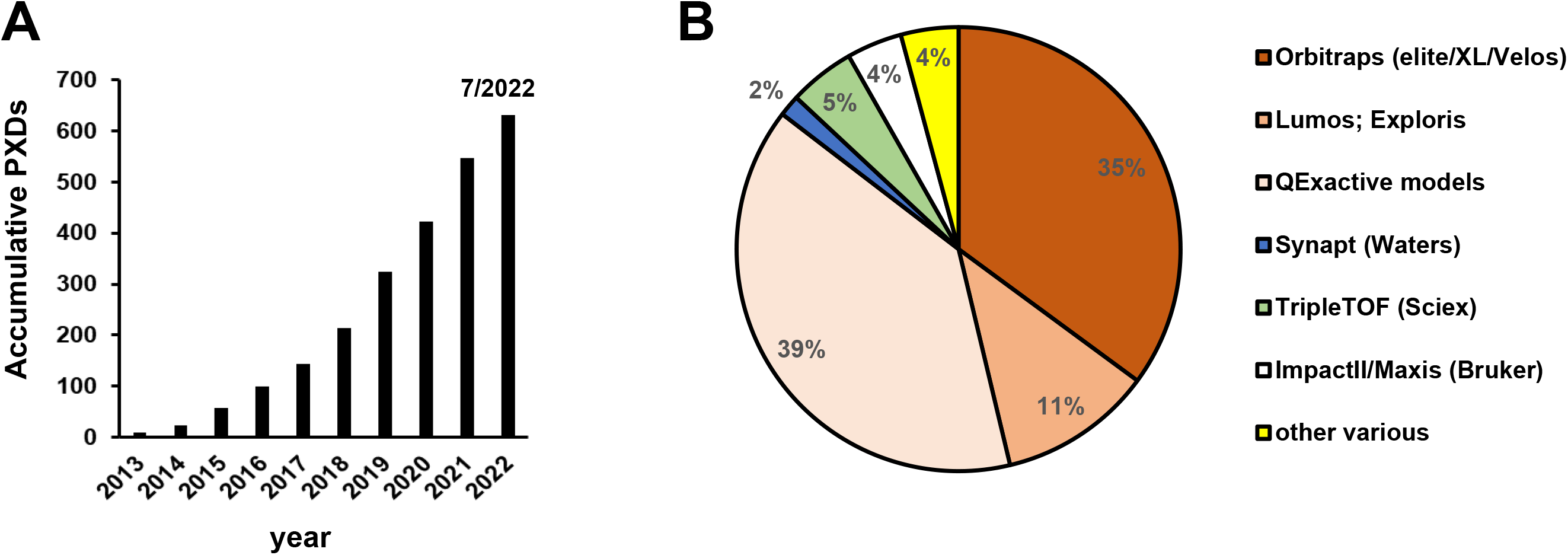
Publicly available PXDs and mass spectrometry instrumentation for *Arabidopsis thaliana* in ProteomeXchange. A, Cumulative PXD available. B, Mass spectrometry instruments used to acquire data in these PXDs (‘other’ includes low resolution instruments such as LCQs, LTQs, QStar, as well as MALDI-TOF-TOF).

**Table 1.**
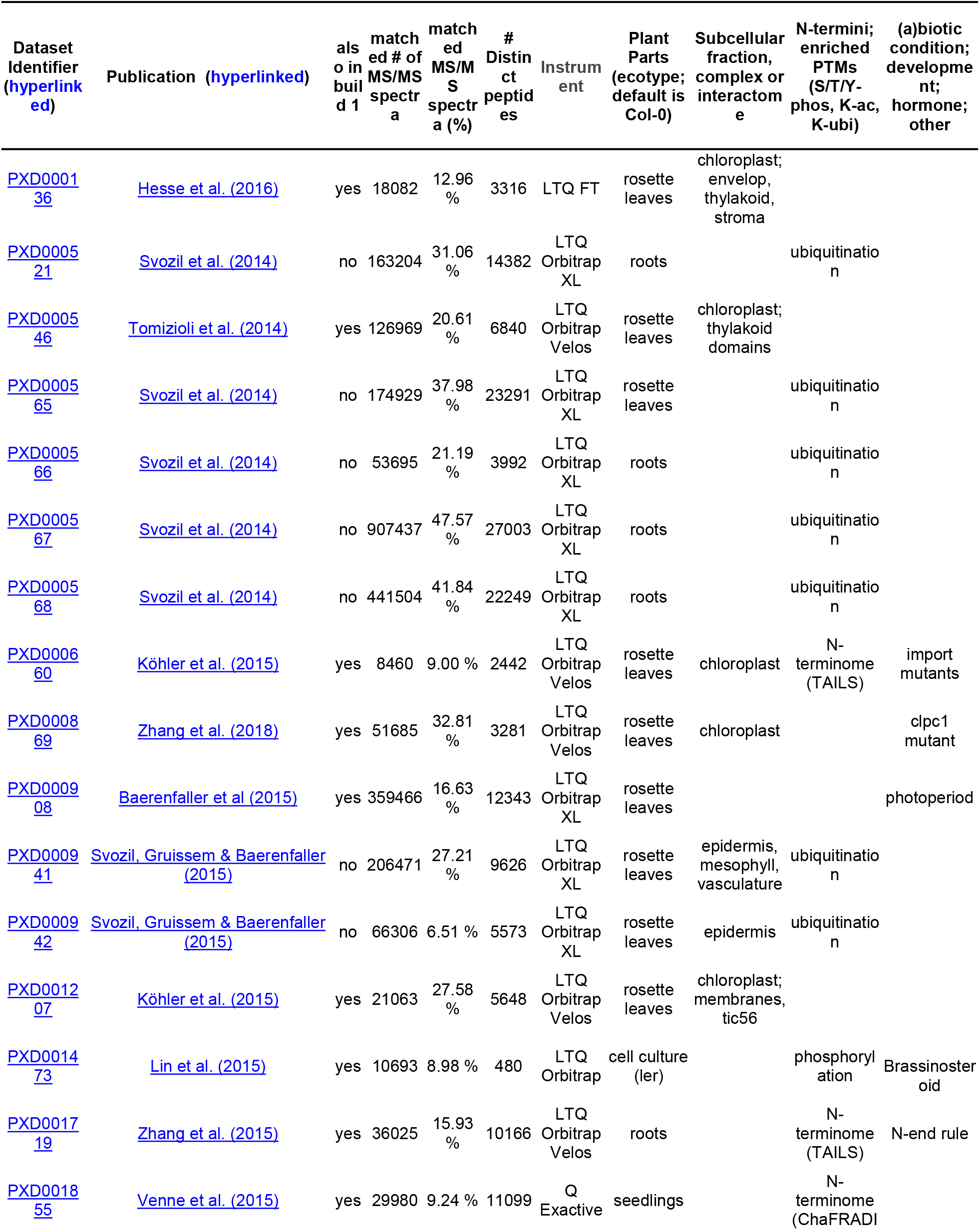
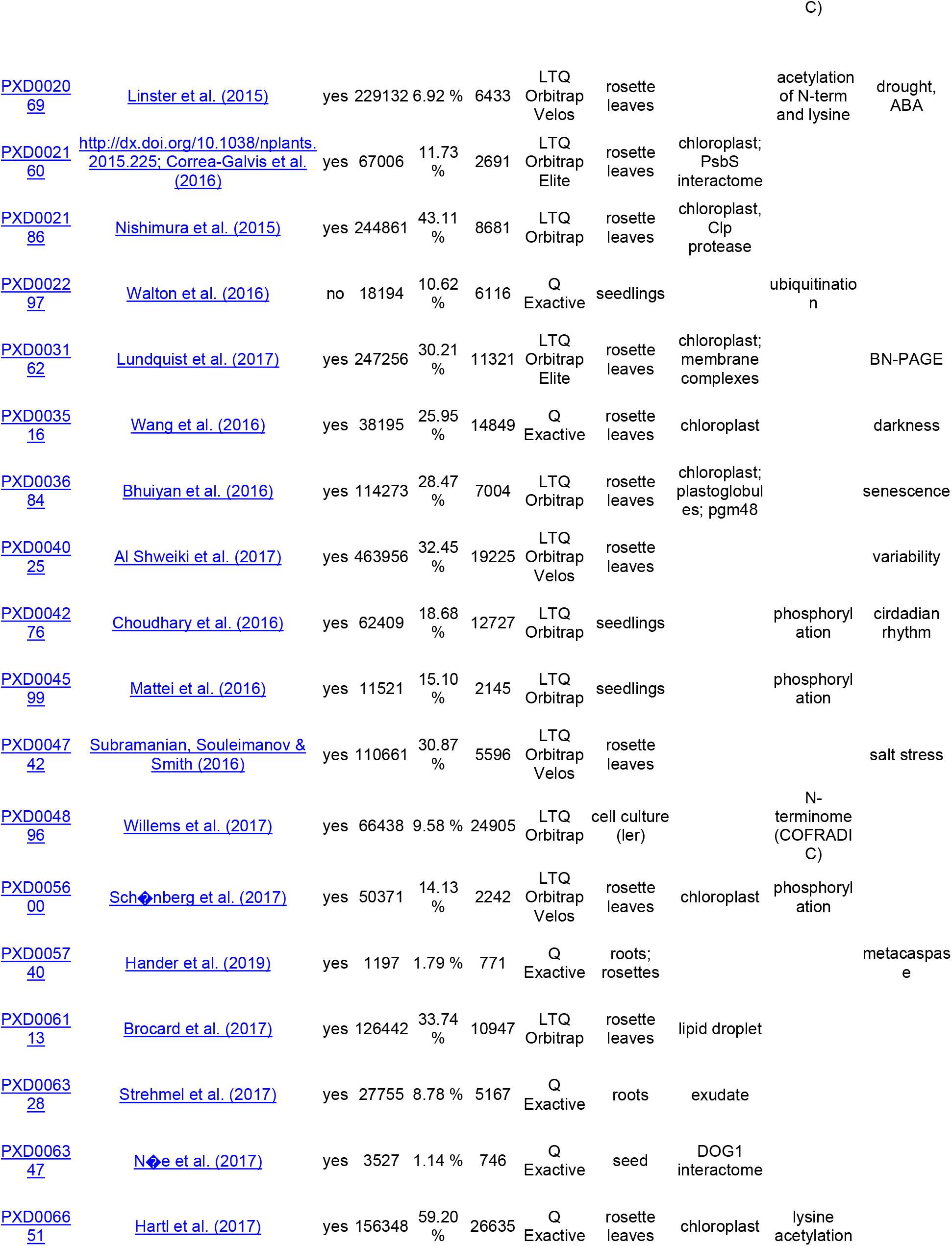
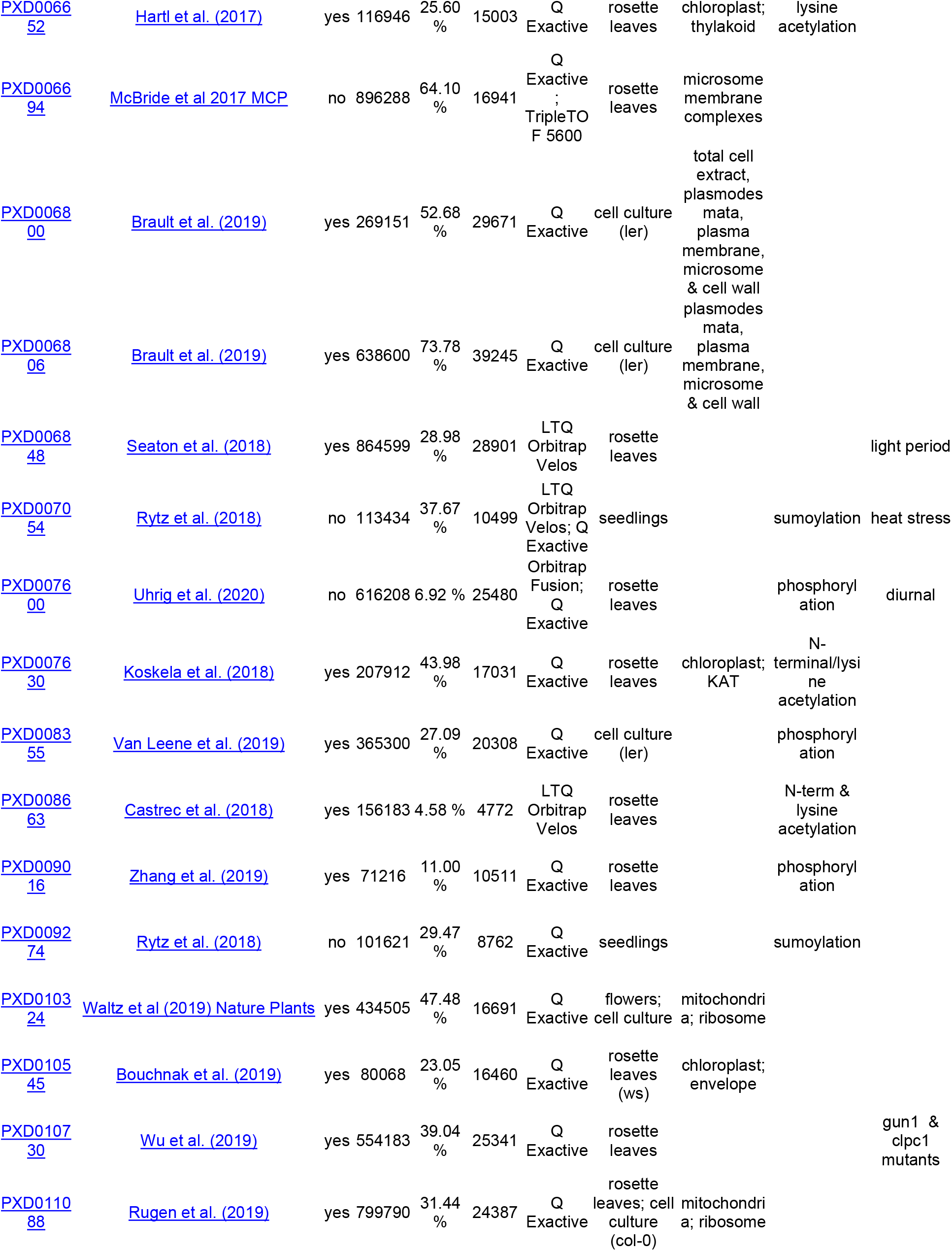
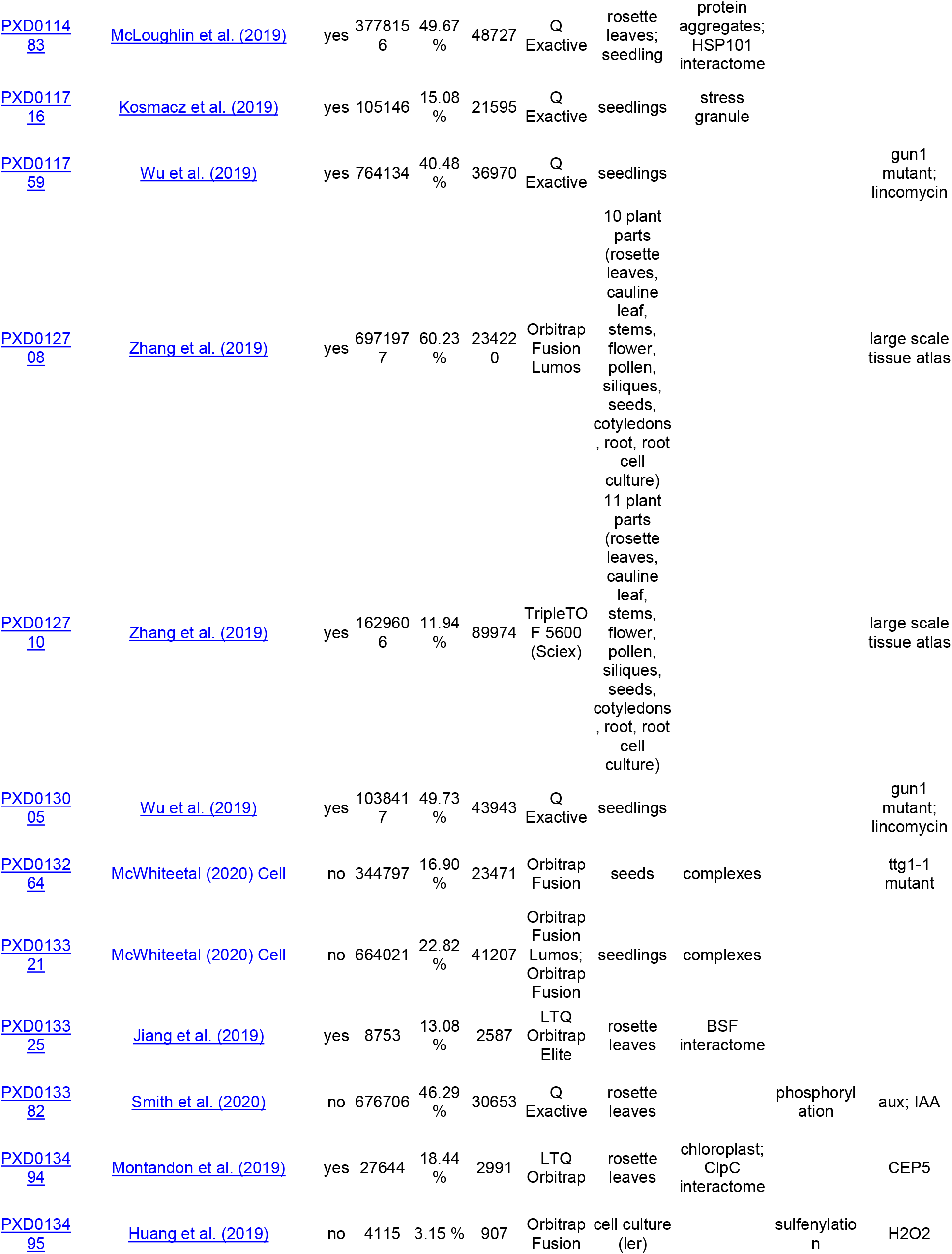
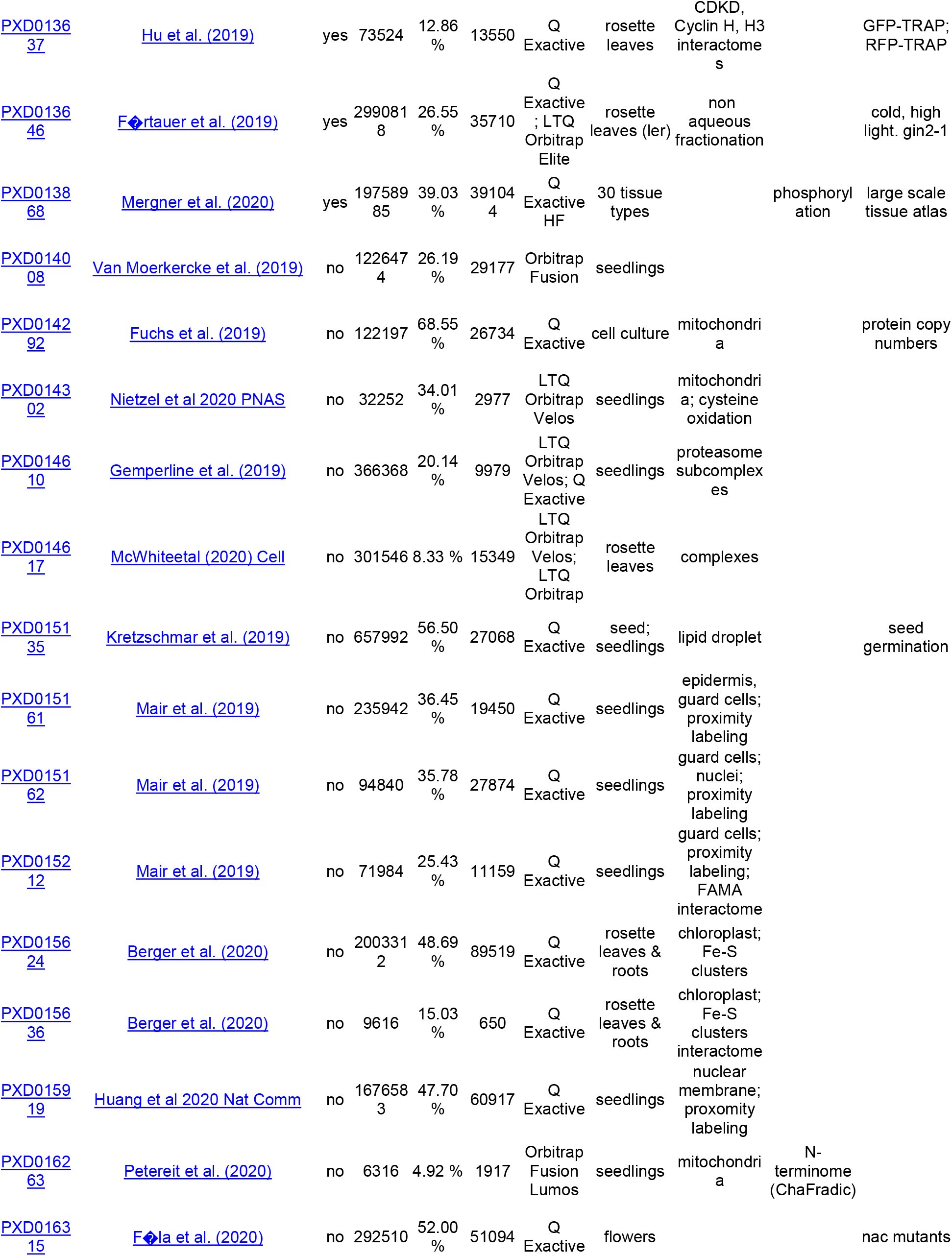
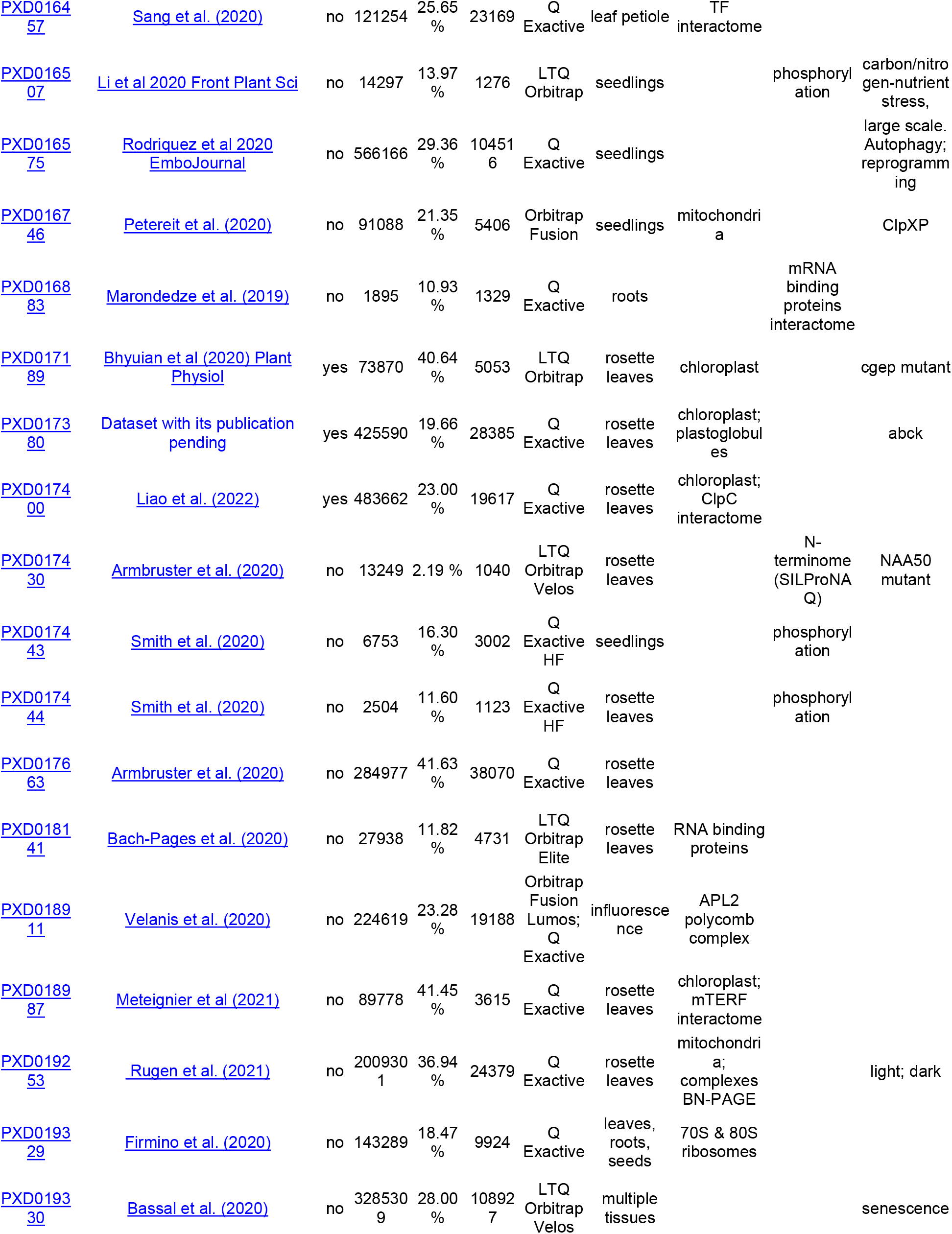
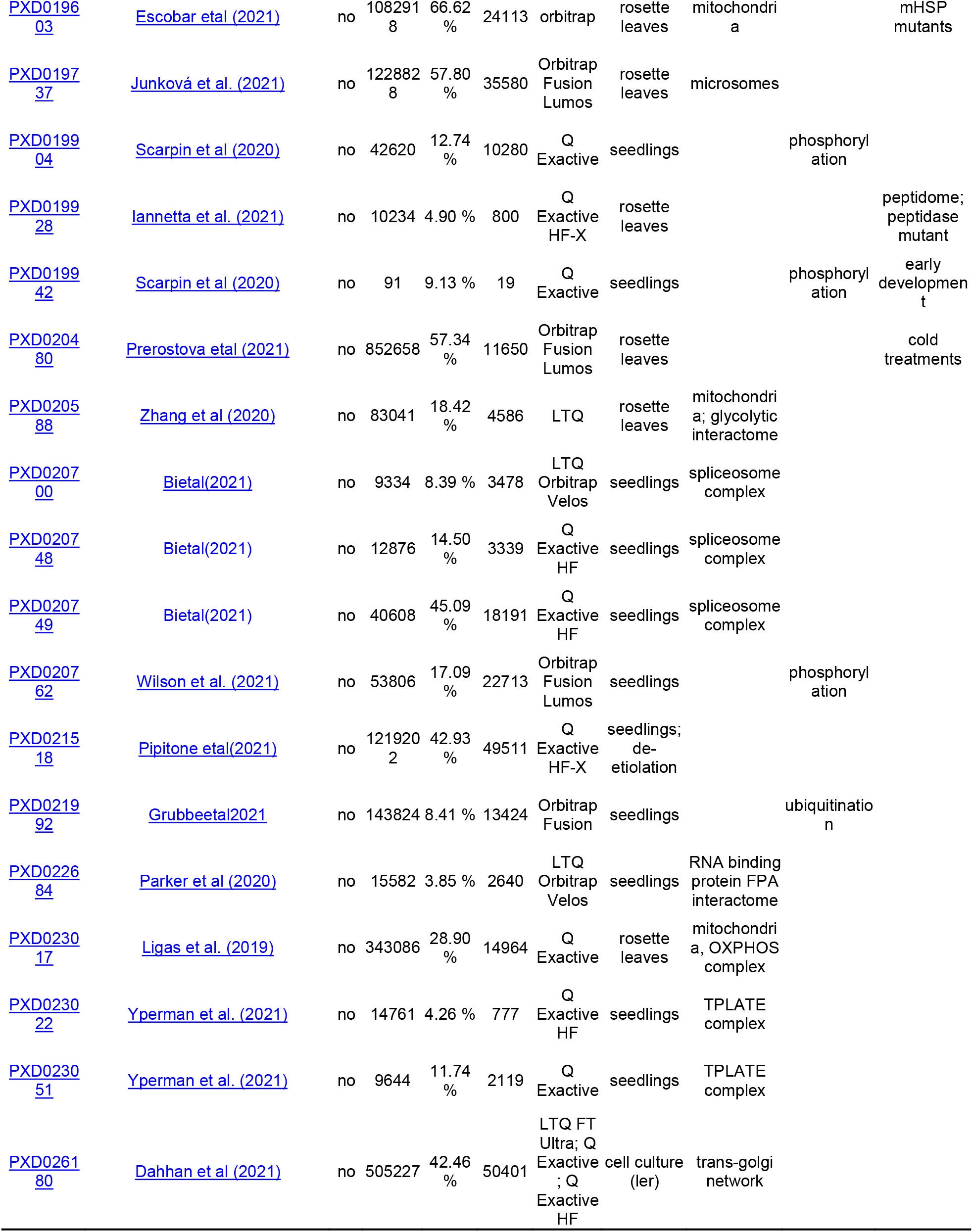
Summarizing information for each PXD in build 2. More details and breakdown into individual experiments are provided in Supplemental Data Set 1 and the metadata annotation system in PeptideAtlas.

### Assembly of a comprehensive protein search space to maximize protein discovery

We assembled a comprehensive protein search space (Table 2) that included the two most recent Arabidopsis annotations (Araport 11 and TAIR10). These are still both used in recent proteomics studies even though Araport11 was released in 2017 (Cheng et al., 2017) and TAIR10 in 2010 (Lamesch et al., 2012). In addition, we added all other Arabidopsis (Col-0) sequences from the universal databases UniProtKB and RefSeq because these are widely used sequence resources. To help identify proteins not represented (or with alternative proteoforms) in these four main resources, we also included sequences generated by individual labs, including a large set of small Open Reading Frames (sORFs) (Hazarika et al., 2017), as well as the predicted protein sequences for 950 Araport11 pseudogenes. These pseudogenes are assumed to be untranslated, but we did previously find evidence that some do appear to produce stable proteins (van Wijk et al., 2021). We also updated the set of the plastid- and mitochondrial-encoded proteins to address redundancies and mistakes in plastid- and mitochondrial ATGC and ATMG sequences, and to allow a systematic analysis of non-synonymous RNA editing for plastid- and mitochondrial encoded proteins. In a forthcoming study (van Wijk et al, in preparation), we will provide detail on the annotation and redundancy of plastid- and mitochondrial encoded proteins, the expression of organellar ORFs, and the impact of RNA editing. Table 2 shows the number of sequences for each sequence data set, their overlap and unique protein sequences.

**Table 2.** Summary of source databases for the Arabidopsis search space.

| Source | Sequences | Distinct | Unique | PeptideAtlasAllOrganellar | PeptideAtlasMinimalOrganellar | AraportUpdated | Araport11 | TAIR10 | Pseudogenes | UniProtKB | RefSeq | ARA-PEP:LW | ARA-PEP:SIPs | ARA-PEP:sORFs | lowaORFs |
| --- | --- | --- | --- | --- | --- | --- | --- | --- | --- | --- | --- | --- | --- | --- | --- |
| PeptideAtlasAllOrganellar (a) | 197 | 195 | 34 |  | 114 | 123 | 106 | 110 | 0 | 110 | 103 | 0 | 0 | 0 | 37 |
| PeptideAtlasMinimalOrganellar (b) | 114 | 114 | 0 |  |  | 114 | 64 | 65 | 0 | 93 | 79 | 0 | 0 | 0 | 27 |
| AraportUpdated (c) | 42617 | 40716 | 0 |  |  |  | 40666 | 31026 | 0 | 38651 | 40660 | 0 | 0 | 0 | 1112 |
| Araport11 (d) | 48359 | 40784 | 10 |  |  |  |  | 31133 | 0 | 38700 | 40654 | 0 | 0 | 0 | 1147 |
| TAIR10 (e) | 35386 | 32785 | 1500 |  |  |  |  |  | 0 | 29401 | 31032 | 0 | 0 | 0 | 1057 |
| Pseudogenes (f) | 3720 | 3702 | 3701 |  |  |  |  |  |  | 0 | 0 | 0 | 0 | 1 | 0 |
| UniProtKB (g) | 39342 | 39273 | 373 |  |  |  |  |  |  |  | 38669 | 0 | 0 | 0 | 1115 |
| RefSeq (h) | 48265 | 40709 | 5 |  |  |  |  |  |  |  |  | 0 | 0 | 0 | 1116 |
| ARA-PEP:LW (i) | 16809 | 16628 | 16478 |  |  |  |  |  |  |  |  |  | 21 | 129 | 0 |
| ARA-PEP:SIPs (j) | 607 | 606 | 565 |  |  |  |  |  |  |  |  |  |  | 20 | 0 |
| ARA-PEP:sORFs (k) | 7901 | 7764 | 7614 |  |  |  |  |  |  |  |  |  |  |  | 0 |
| lowaORFs (l) | 7481 | 7270 | 6116 |  |  |  |  |  |  |  |  |  |  |  |  |
| **Total non-redundant** |  |  |  |  |  |  |  |  |  |  |  |  |  |  |  |

(a) PeptideAtlasAllOrganellar includes all the PeptideAtlas\_ATxGnnnnnn.1 (original), .2 (RNA edits [major]), .3 (RNA edits [major and minor]), .4 (RNA edits [major, minor, and truncations])
(b) PeptideAtlasMinimalOrganellar includes one protein for each organellar gene, the RNA edited [major only] version if there are edits, or the original if no editing sites
(c) AraportUpdated begins with the Araport11 proteome with all organellar proteins replaced with PeptideAtlasMinimalOrganellar set and other corrections discussed in this article applied
(d) Araport11 represents the current set of Araport11 proteins as downloaded 2021-04-26
(e) TAIR10 represents the current set of TAIR10 proteins as downloaded 2020-12-22
(f) Pseudogenes represent an additional set of entries labeled as "pseudogenes" in Araport11 and are thus not exported as part of the proteome - downloaded '21-02-23

### Protein identification, sequence coverage, PTMs and overall statistics in build 2

The 115 selected PXDs contained 259.4 million raw MSMS spectra from 10478 MS runs that we searched as 369 different experiments (Tables 3 and Supplemental Data Set S1). We assigned these experiments based on the metadata associated with the PXDs, as well as associated publications. Importantly, this involved manual evaluation of experimental conditions, sample preparations and proteomics and MS workflows; this is a relatively time-consuming process requiring specific expertise which is currently hard to automate. This allowed us to search with appropriate parameters (parameters need to be assigned for specific PTMs, protease cleavage reagents, iTRAQ, TAILS, COFRADIC) and also to associate the most relevant biological (*e.g.* dark vs light treatments) and technical metadata. The associated metadata will facilitate discoveries of biological relevance (*e.g.* condition or cell-type specific accumulation patterns, the relation between alternative splicing and plant material), but also to analyze for technical features (*e.g*. sample-handling related PTMs such as off-target effects of iodoacetamide (Hains and Robinson, 2017; Muller and Winter, 2017) or trypsin artefacts (Schittmayer et al., 2016; Niu et al., 2020)

In total there were 70.5 million peptide-spectrum matches (PSMs) to nearly 0.6 million distinct peptides, thereby identifying 18267 Araport11 proteins at the highest confidence level (canonical proteins, two uniquely mapping non-nested peptides of ≥ 9 residues and with ≥ 18 residues of total coverage) and 1856 ‘uncertain’ proteins (too few uniquely-mapping peptides of ≥ 9 aa to qualify for canonical status and may also have one or more shared peptides with other proteins) and 1540 ‘redundant’ proteins (containing only peptides that can be better assigned to other entries and thus these proteins are not needed to explain the observed peptide evidence) (Table 3). The overall FDR of the PSMs was 0.08%. The ‘uncertain’ proteins are needed to explain all the peptides identified above threshold, while ‘redundant’ identifications have only peptides that already map to canonical or uncertain proteins – for more details on these definitions see (van Wijk et al., 2021). These ‘redundant’ proteins typically have significant sequence homology to these canonical proteins. Table 4 shows the breakdown of identifications at different confidence levels for each of the five nuclear chromosomes, as well as the plastid and mitochondrial genomes. The percentage of identified predicted proteins per nuclear chromosome was on average 78.6% with only small differences between chromosomes. We identified nearly all predicted mitochondrial and plastid proteins and ORFs (91% and 95%, respectively); the few unobserved organellar proteins are either untranslated ORFs (likely pseudogenes) or very small proteins. In summary, build 2 has peptides mapping to 78.6% (21663/27559) of all predicted proteins in Araport11 (counting only one isoform per gene). The complete sets of identified proteins in their respective confidence tiers can be downloaded at https://peptideatlas.org/builds/arabidopsis/

**Table 3.** Comparison of summary statistics of Arabidopsis PeptideAtlas Builds 1 and 2.

| Metric | Build 1 | Build 2 | Ratio of 2 / 1 |
| --- | --- | --- | --- |
| Datasets (PXDs) | 52 | 115 | 2.21 |
| Experiments | 266 | 369 | 1.39 |
| MS Runs | 6,148 | 10,478 | 1.70 |
| MS2 Spectra Acquired (a) | 142,703,610 | 259,383,093 | 1.82 |
| MS2 Spectra Scored (b) | 125,181,633 | 210,655,824 | 1.68 |
| PSM FDR | 0.001 | 0.0008 | 0.80 |
| PSMs passing threshold | 39,480,811 | 70,470,125 | 1.78 |
| Distinct Peptides | 535,340 | 596,839 | 1.11 |
| Canonical proteins (Araport11*) | 17,858 | 18,267 | 1.02 |
| Uncertain proteins (Araport11*) | 1,942 | 1,856 | 0.96 |
| Redundant proteins (Araport11*) | 1,600 | 1,540 | 0.96 |
| Not observed proteins (Araport11*) | 6,255 | 5,896 | 0.94 |
| Araport11* proteins with peptides mapped | 21,400 | 21,663 | 1.01 |
| (a) information in raw files |  |  |  |
| (b) spectra of sufficient quality to be scored |  |  |  |
| * Araport11 but with updated plastid and mitochondrial encoded proteins (114 instead of 210 in original Araport11) and total size is 27559 proteins |  |  |  |

**Table 4.** Proteins identified in Araport11* for each of the four confidence categories in build 2 for mitochondrial-(M) and plastid (C) chromosomes and the nuclear chromosomes (1-5).

| Chromosome* | Entries | Canonical, n (%) |  | Uncertain, n (%) |  | Redundant , n (%) |  | Not Observed , n (%) |  |
| --- | --- | --- | --- | --- | --- | --- | --- | --- | --- |
| **M** | 35 | 27 | 77.1% | 5 | 14.3% | 0 | 0.0% | 3 | 8.6% |
| **C** | 79 | 63 | 79.7% | 12 | 15.2% | 0 | 0.0% | 4 | 5.1% |
| **1** | 7156 | 4730 | 66.1% | 502 | 7.0% | 384 | 5.4% | 1540 | 21.5% |
| **2** | 4317 | 2762 | 64.0% | 290 | 6.7% | 240 | 5.6% | 1025 | 23.7% |
| **3** | 5460 | 3630 | 66.5% | 353 | 6.5% | 296 | 5.4% | 1181 | 21.6% |
| **4** | 4180 | 2788 | 66.7% | 282 | 6.7% | 247 | 5.9% | 863 | 20.6% |
| **5** | 6332 | 4267 | 67.4% | 412 | 6.5% | 373 | 5.9% | 1280 | 20.2% |
| **Total** | 27559 | 18267 | 66.3% | 1856 | 6.7% | 1540 | 5.6% | 5896 | 21.4% |

\* Araport11 but with updated plastid and mitochondrial encoded proteins (114 instead of 210 in original Araport11) and total size is 27559 proteins

In addition, there were 4342 peptides only matching to proteins in sources other than Araport11 with a total of 1.8 million PSMs (Table 5). These peptides were assigned to proteins by hierarchy of sources (ranked from 1 to 11), with each peptide assigned only to the highest-ranking source possible and then not to any other source. Table 5 also summarizes how many of these non-Araport11 proteins were identified when applying different thresholds for the minimum number of PSMs and matched peptides. For example, when requiring at least 2 distinct peptides with each at least 3 observations (PSMs) there are 25 proteins identified in TAIR10 and nine pseudogenes, as well as 6 small proteins (LW or sORFs) from the ARA-PEP database. Supplemental Data Set S3 provides more information on these proteins not found in Araport11. These matched pseudogenes and non-Araport11 proteins should be considered for incorporation into the next Arabidopsis genome annotation. Finally, what this Table 5 also demonstrates is that samples also contain various contaminants (*e.g*. keratins from human skin, trypsin for auto-digestion, BSA), as expected based on observations in other large-scale studies (Hodge et al., 2013; Frankenfield et al., 2022).

**Table 5.** Peptides assigned to proteins by hierarchy of sources ranging from Araport11 to DECOY, with each peptide is assigned only to the highest source possible and then not to any other source.

| Hierarchy (a) | Primary Protein Match | No. of peptides | No. of PSMs | No. of Primary Proteins | No. of peptides ($\geq 3$ PSMs) | No. of PSMs ($\geq 3$ PSMs) | No. of Primary Proteins ($\geq 3$ PSMs) | No. of Primary Proteins ($\geq 2$ Distinct Peptides with $\geq 3$ PSMs) |
| --- | --- | --- | --- | --- | --- | --- | --- | --- |
| 1 | Araport11 | 595346 | 70409850 | 20876 | 411364 | 70166505 | 18860 | 17056 |
| 2 | TAIR10 | 438 | 33123 | 69 | 271 | 32908 | 43 | 25 |
| 3 | PSEUDOGENE | 205 | 1264 | 126 | 74 | 1104 | 54 | 9 |
| 4 | UniProtKB | 197 | 14408 | 38 | 120 | 14306 | 28 | 15 |
| 5 | RefSeq | 0 | 0 | 0 | 0 | 0 | 0 | 0 |
| 6 | ARA-PEP:LW | 101 | 519 | 82 | 30 | 440 | 21 | 3 |
| 7 | ARA-PEP:SIPs | 5 | 17 | 5 | 2 | 14 | 2 | 0 |
| 8 | ARA-PEP:sORFs | 75 | 404 | 60 | 29 | 352 | 18 | 3 |
| 9 | IowaORFs | 466 | 10409 | 157 | 232 | 10111 | 61 | 26 |
| 10 | CONTAM(b) | 5217 | 1719466 | 95 | 3577 | 1717256 | 88 | 83 |
| 11 | DECOY(c) | 728 | 8001 | 654 | 281 | 7470 | 269 | 10 |
| (a) Hierarchy refers to the order to which peptides are assigned to sources. |  |  |  |  |  |  |  |  |
| (b) Contaminants often found in samples, e.g. BSA, Keratin, trypsin, etc |  |  |  |  |  |  |  |  |
| (c) Decoys are all shuffled protein sequences in the search space; this enables accurate calculation of FDR. |  |  |  |  |  |  |  |  |

Build 2 contains more than double the number of PXDs as build 1, and 68% more raw MSMS spectra were searched (Table 3). Whereas the number of PSMs increased by 78%, the number of distinct identified peptides only increased by 11% and the number of identified proteins (across all confidence levels) increased by just 1% (Table 3). Figure 2A shows the cumulative identified peptides as well as distinct peptides from the 369 experiments (each PXD can have more than one experiment), whereas figure 2B shows the cumulative identified canonical proteins as well as distinct canonical proteins from the experiments. This shows that even though we deliberately selected PXDs to enrich for underrepresented proteins, this did only incrementally increase peptide and protein discoveries, despite the near doubling of matched PSMs. This clearly suggest that identification of the remaining 21% of the predicted proteome will require new approaches.

**Figure 2.**
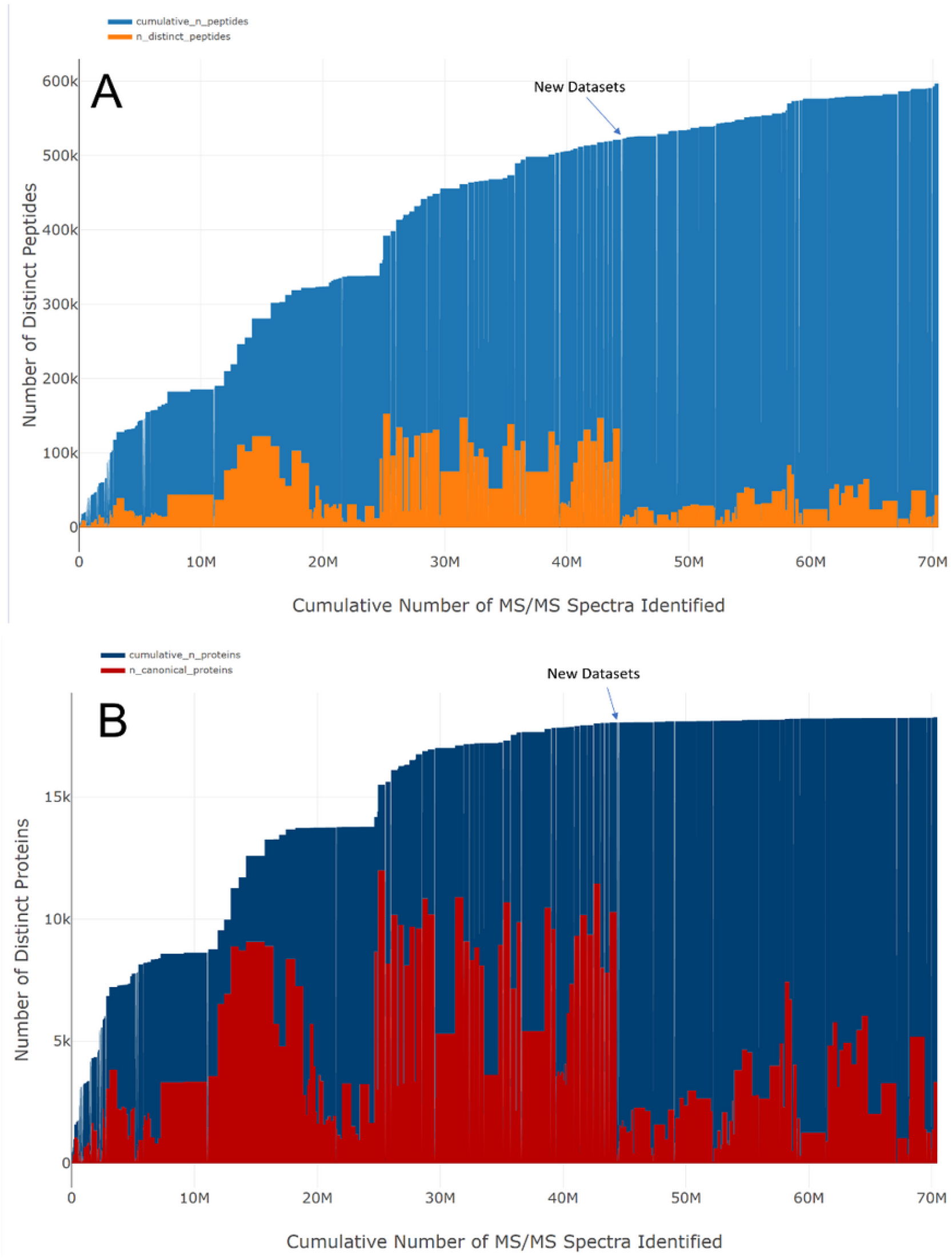
Contributions of individual experiments to the PeptideAtlas Build. A, From the 369 experiments conducted, the graph displays the total number of distinct peptides for the build as well as the number of peptides contributed by each experiment. B, The plot shows the cumulative number of distinct proteins and the number of proteins that were contributed from each experiment. The location where new datasets added since the first build is marked.

To better understand possible underlying causes for these diminished returns, we investigated the relationships between number of matched spectra and identified distinct peptides or proteins for each PXD. This showed a wide PSM match rate for searched spectra between PXDs ranging from 1% to 74% (Table 1) mostly due to differences in spectral quality (due to *e.g.* peptide abundance, instrument settings and sensitivities, sample preparation), but a strong positive linear correlation between the number of matched spectra and identified distinct peptides or distinct proteins (Supplemental Figures S1,2). Interestingly, plotting the % of matched spectra to identified distinct peptides or proteins showed a clear saturation (or diminished return) suggesting bottlenecks in the dynamic range for protein identification (Supplemental Figures S1,2). This suggests that dramatic innovations in mass spectrometry and/or proteomics workflows and sample selection are needed to identify the remaining 21.4% of the predicted proteome.

### Mapping biological PTMs; N-terminal and lysine acetylation, phosphorylation and ubiquitination

We selected multiple PXDs that specifically enriched for the physiologically important PTMs of phosphorylation, N-terminal acetylation, lysine acetylation or ubiquitination (Table 1). A sophisticated PTM viewer in PeptideAtlas allows detailed examination of these PTMs, including direct links to all spectral matches. PTM identification rates strongly depend on the confidence level (minimal probability threshold) of PTM assignment. We limited our summary in this publication on PTMs to canonical proteins, but PTMs for all confidence levels of protein identification are available in the PeptideAtlas web interface. Here we used localization probability P≥0.95 from PTMProphet (Shteynberg et al., 2019) for each PTM, and also required at least 3 PSMs for a specific PTM at a specific residue to be included in the overall statistics. In general, higher numbers of repeat observations (PSMs) for a specific PTM at a residue improve the reliability of the assignment. Conversely, peptides with high PSM counts (*e.g.* hundreds or more) for which the vast majority (*e.g.* 99%) of peptide do not have a reported PTM at P>0.95, are possibly false discoveries. We recommend therefore to use the PeptideAtlas to evaluate specific PTM sites if these are of particular interest to the reader. We evaluated the results for false positives and possible pitfalls in various ways, including spot checking matched spectra and proteins to which PTMs were mapped. Supplemental Data Sets S4-S7 provide the results for these four PTMs and Supplemental Data Set S8 provides the combined results of these PTMs per canonical protein to analyze for possible cross-talk between PTMs. We briefly summarize the results below:

### N-terminal acetylation (NTA)

Proteins are synthesized with an initiating N-terminal methionine which can be N-terminally acetylated. However, a large portion of cellular proteins undergo removal of the initiating methionine residue by methionine amino peptidases (MAPs) if the side chain of the second residue is small enough (Giglione et al., 2004; Ross et al., 2005). If the N-terminal methionine is removed, NTA can occur on the 2nd residue of the predicted protein. Both methionine removal and NTA are co-translational processes that occur in the cytosol and plastids (Willems et al., 2021; Meinnel and Giglione, 2022; Pozoga et al., 2022). However, nuclear-encoded proteins synthesized in the cytosol and then sorted into chloroplasts, can undergo post-translational NTA after removal of the cleavable chloroplast transit peptide (cTP) by several N-terminal acetyltransferases (NATs) in the chloroplast (Meinnel and Giglione, 2022; Pozoga et al., 2022) Indeed, intra-chloroplast NTA has been documented by several studies mostly involving N-terminal labeling with stable isotopes followed by fractionation (TAILS, SILProNAQ, COFRADIC) (Dinh et al., 2015; Rowland et al., 2015; Bienvenut et al., 2020; Willems et al., 2021) and won’t be further addressed in this study. The presence of NATs in the nucleus (NAA50), ER (NAA50) and plasma membrane (NAA60) allows for additional post-translational NTA after sorting to these respective subcellular locations (Pozoga et al., 2022), thus adding to the complexity of NTA patterns. Finally, proteins sorted to mitochondria with cleavable N-terminal sorting signals typically do not accumulate with an acetylated N-terminus (Huang et al., 2009) and indeed no NAT has been reported to localize to mitochondria. When peptides are identified matching to the initiating methionine or the immediate downstream residue of a protein, this is important support for the lack of cleavable N-terminal sorting signals (because the sorting and cleavage process and subsequent degradation of the cleaved signal peptide is typically very efficient).

After removal of false positives (see below), the search process identified 3185 Araport11 canonical proteins (including 18 chloroplast- and 5 mitochondrial-encoded proteins) containing 3258 NTA sites mostly at position 1 (M) or position 2, and the remainder further downstream (Supplemental Data Set S4). 98% of these NTA proteins contained a single NTA site. The 2% of cases where more than one NTA site per protein was found could be due to alternative splice forms or translation start sites (Willems et al., 2021), proteins sorted to one or more subcellular location or false discovery of the PTM (there is no known sample preparation induced NTA). Interestingly, we found 30 false positive NTAs in four (iso)leucine-repeat peptides (sequences: IIIIIIIIII or VIIIIII or VIIIIIII or VVLLIIL matching to 27 canonical proteins). [Acetyl]-V has an identical mass as [Formyl]-L or I (L and I are isobaric) and these false positives stem from this misassignment. Formylation can occur at any peptide N-terminus (and the side chain of T and S) and is a common PTM induced by formic acid (even at low concentrations) (Zybailov et al., 2009; Kim et al., 2016). We also noted false positives due to combinatorial (assigned or real) mass modifications, involving deamidation (+0.98402 Da), carbamylation (+43.00582 Da) and C12/C13 isotopes (+1 Da), especially when the assigned NTA (+42.01056 Da) was observed with an absolute low number of PSMs or a relative low number of PSMs compared to the total number of PSMs for that peptide (for highly abundant proteins).

There were 1493 nuclear-encoded canonical proteins with matched peptides starting exclusively with the initiating methionine, of which 1164 were observed with NTA. There were 2810 nuclear-encoded canonical proteins with matched peptides starting exclusively at position 2, of which 1912 were observed with NTA. These acetylated residues were mostly for proteins without predicted N-terminal signal peptides (sP, cTP or mTP). We created sequence logo plots for each of these four groups (Figure 3A-D) to show the methionine amino peptidase activity (to remove the initiating M) and the NAT activity. The logos show that proteins that retain the methionine have mostly the acidic amino acids residues (D,E) and N in the 2nd position (Figure 3A). NTA occurs on the initiating M (Figure 3C), as well as on A,S,V,G (Figure 3D). The iceLogo (Maddelein et al., 2015) (Figure 3E) comparing the sets in panel B and D shows that the dominant NAT activity for this set of identified proteins is to acetylate A and S residues. NTA is the result of the activity of multiple NATs each with their own set of preferred substrates and NATA has been reported to be responsible for N-terminal acetylation of ∼50% of the plant proteome (Pozoga et al., 2022).

**Figure 3.**
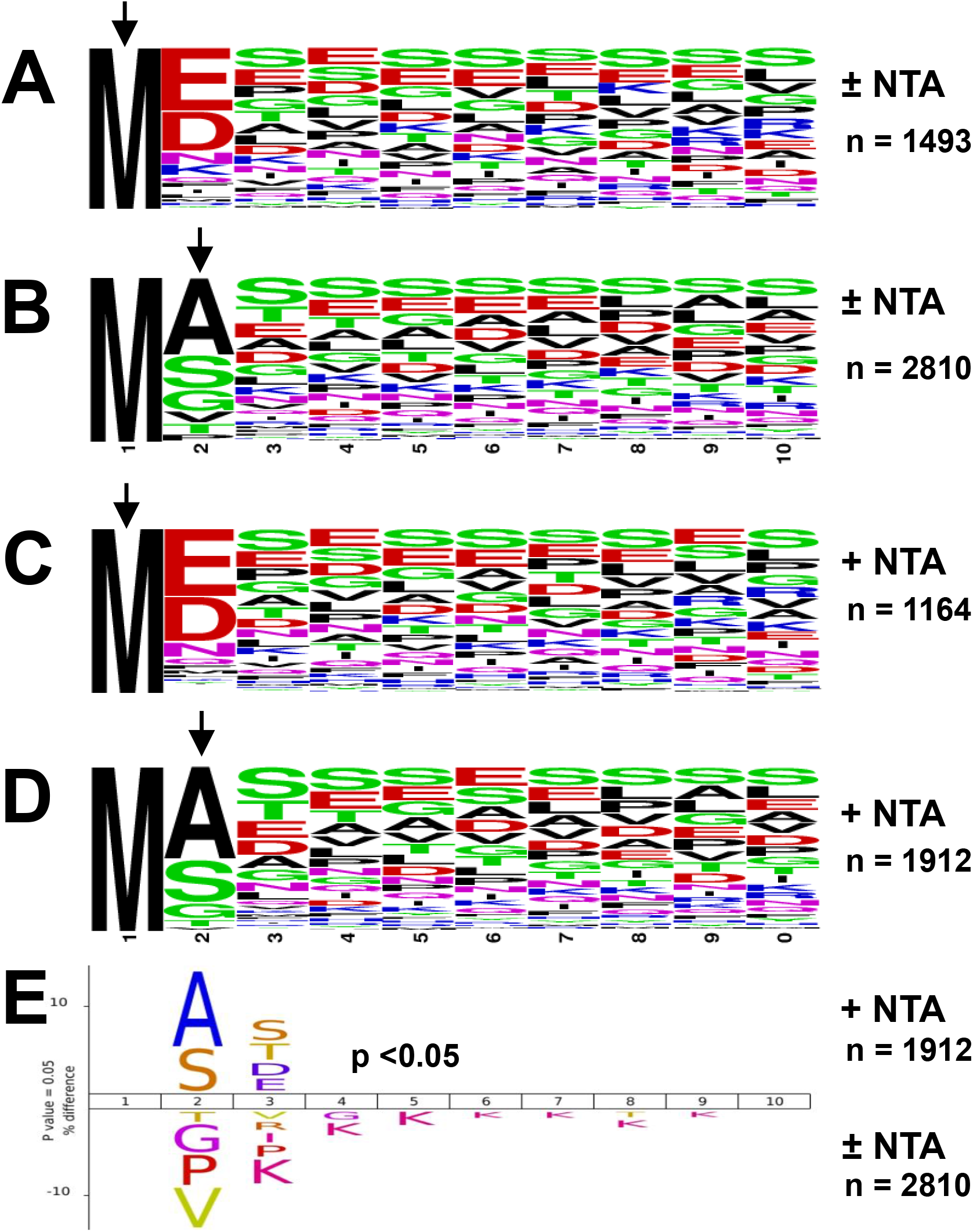
N-terminal consensus sequence patterns of canonical nuclear-encoded proteins accumulating with the initiating methionine or the 2^nd^ residue (after methionine excision) with or without NTA. A,B, Sequence logos of proteins (first 10 residues are shown) that are exclusively found with the initiating methionine (A) or exclusively found with just this methionine removed (B), irrespective of NTA. C,D, Sequence logos of NTA proteins (first 10 residues are shown) exclusively accumulating with the initiating methionine (C) or exclusively found with the second residue (methionine removed). E. Icelogo for NTA canonical proteins exclusively starting at position 2 using all canonical protein starting exclusively at position 2, but irrespective of the NTA status. Arrows indicate the observed N-terminal residue.

### Lysine acetylation

Identification of K-acetylation required a targeted search that was applied on the raw files from three PXDs with enriched lysine acetylome samples from the Finkemeier lab (PXD006651, PXD006652, PXD007630) (Table 1). After application of our post-search selection criteria (PTM localization P > 0.95; ≥3 PSMs per PTM site) and removal of false positives, we identified 864 core canonical proteins containing K-Acetyl modifications representing 1750 K-sites (Supplemental Data Set S5). 512 proteins (59%) contained a single K-acetyl site whereas others are more heavily K-acetylated. The acetylated proteins were distributed across multiple subcellular locations and functions supporting recent findings in Arabidopsis (Tilak et al., 2023), but also other plant species (Zhang et al., 2022), the green algae *Chlamydomonas reinhardtii* (Fussl et al., 2022) as well as the moss *Physcomitrium patens* (Balparda et al., 2022)

### Phosphorylation

After application of our post-search selection criteria (PTM localization score P>0.95; ≥3 PSMs per PTM site), there are 5198 canonical phosphoproteins (p-proteins) representing 14748 phosphosites (p-sites) (86% S, 13% T, 0.6% Y) (Supplemental Data Set S6). 45% of the 5198 p-proteins contained only a single p-site, and 20%, 11% and 7% contained 2, 3 or 4 p-sites, respectively. This ratio between pS, pT and pY is consistent with published literature for large scale phosphorylation data sets in Arabidopsis (van Wijk et al., 2014; Mergner et al., 2020).

### Ubiquitination

We found 668 ubiquitinated core canonical proteins based on 765 single K-glycine (KG) sites (Walton et al., 2016) and 412 K-diglycine (KGG) sites (Grubb et al., 2021), totaling 1177 ubi-sites (Supplemental Data Set S7). The two PXDs that contained enriched ubiquitinated sites were from large scale studies (Walton et al., 2016; Grubb et al., 2021) that applied different methods (resulting in K-G or K-GG) to identify the ubiquitinated sites. 449 proteins (67%) contained a single G or GG PTM site. By far the most PSMs for G or GG were found for nine ubiquitin (extension) proteins (>1000 PSMs), followed (albeit at far lower PSM levels) by several plasma membrane proteins and histones. We note that there are no mitochondrial-encoded proteins and one chloroplast-encoded protein PeptideAtlas_ATCG00900.1 (30S ribosomal protein S7A/B) with just three PSMs for one site (K13-G). 45 sites across 18 proteins exhibited both a Gly and a GG PTM. Since the G and GG studies were independent, this might indicate that these sites have a lower FDR than sites which were only detected by one of the methods. These 18 proteins are the nine ubiquitin or ubiquitin extension proteins which is logical since they form polyubiquitination chains. The others are abundant glycolytic enzymes (aldolases), cytosolic ribosomal proteins, an elongation factor involved in cold-induced translation (LOS1)(Guo et al., 2002), the SNARE protein AtVAM3p (Sanderfoot et al., 1999), and two enzymes involved in amino acid metabolism (Supplemental Data Set S7). It is perhaps not surprising that there was so little overlap between ubiquitination sites between these two studies because ubiquitination is generally a transient PTM, and in case of polyubiquitination this leads to rapid degradation. Furthermore, plant materials, sampling and methodologies were very different across these two studies.

### Summary of the PTMs

All together, we identified 5764 proteins with one or more of these four PTMs (NTA, Kac, P, or UBI) based on 0.582 million PSMs for 17675 PTM sites (Supplemental Data Set S8). 4952 proteins contain only one type of PTM, 635 proteins contain two types of PTMs, 160 proteins contain three types of PTMs, and 17 proteins contain all four types of PTMs.

In addition to these physiological PTMs (which require specific affinity enrichment, except for N-terminal acetylation), the MS searches also include additional mass modifications, many of which are induced during sample processing (see Materials and Methods). The frequencies of these can greatly vary between PXDs and experiments within PXDs depending on the use of organic solvents, urea, oxidizing conditions, temperature, pH and use of SDS-PAGE gels. These mass modifications are included in the search parameters since many of these modified peptides would otherwise not be identified or lead to false assignments. However, we do not report on these statistics as they have generally very little physiological relevance. These mass modifications are all available in the PeptideAtlas web interface with viewable spectra and they can be investigated to better understand the impact of different sample treatments.

### Understanding the nature of the unobserved proteomes in the new release of Arabidopsis PeptideAtlas

Of the 27559 predicted nuclear and organelle protein coding genes of the Arabidopsis in Araport11, we identified 18267 (66.3%) corresponding proteins as meeting the canonical criteria (canonical proteins) and 5896 proteins (21.3%) having no observations at all (dark proteins) in our PeptideAtlas build. The remaining identified proteins are in the uncertain or redundant categories. Our working hypotheses is that the dark proteins are not observed because they: i) are generally expressed at too low levels for detection, ii) are expressed only under very specific conditions or in specific cell types, iii) have very short half-life, iv) have physicochemical properties (very small and/or very hydrophobic) that make them difficult to detect using standard proteomics and mass spectrometry workflows (van Wijk et al., 2021), or v) simply not translated at all under any conditions.

Figure 4A displays the histograms of molecular weight (between 0 and 80 kDa) for the canonical and dark proteins. Figure 4B displays the relative proportion of canonical and dark proteins in each kDa bin. Below 4 kDa all proteins are dark proteins. Between 14 and 16 kDa, ∼50% of the proteins are canonical and ∼50% are dark. With increasing molecular weight, the proportion that are canonical proteins increases to ∼90%. There are a substantial number of proteins above 80 kDa, but the proportion of proteins that are canonical is generally constant above 80 kDa at ∼95%. Figure 4C,D displays the distribution of hydrophobicity computed as the gravy score based on the algorithm of Kyte and Dolittle (Kyte and Doolittle, 1982). Values above 0 are considered hydrophobic, with values above 0.5 being very hydrophobic. Figure 4C shows the absolute number of proteins per bin, whereas panel 3D shows the relative proportion of canonical and dark proteins per bin. The two distributions are broadly similar between gravy scores -2.0 to +0.8, with a sharp decline in the proportion of canonical proteins above a gravy score of +0.8. All 64 proteins with a gravy score greater than +1.0 are dark (*i.e.* undetected) and most of these proteins are small with a predicted signal peptide for secretion to the ER. Furthermore, most have no known function, but also include seven arabinogalactan proteins (AGPs) (Silva et al., 2020) and four plasma membrane RCI2 proteins (Medina et al., 2007). Figure 4E,F displays the distribution of isoelectric point (pI) for proteins. Both canonical and dark proteins exhibit the typical bimodal distributions peaking at just below 6.0 and again just above 9.0 based on their total counts (Figure 4E). The distribution in the relative proportion of canonical to dark proteins is complex (Figure 4F), but in general, the proportion of canonical proteins is substantially reduced at the two extremes. Very basic proteins (pI) are enriched for ribosomal proteins and ‘hypothetical’ proteins.

**Figure 4.**
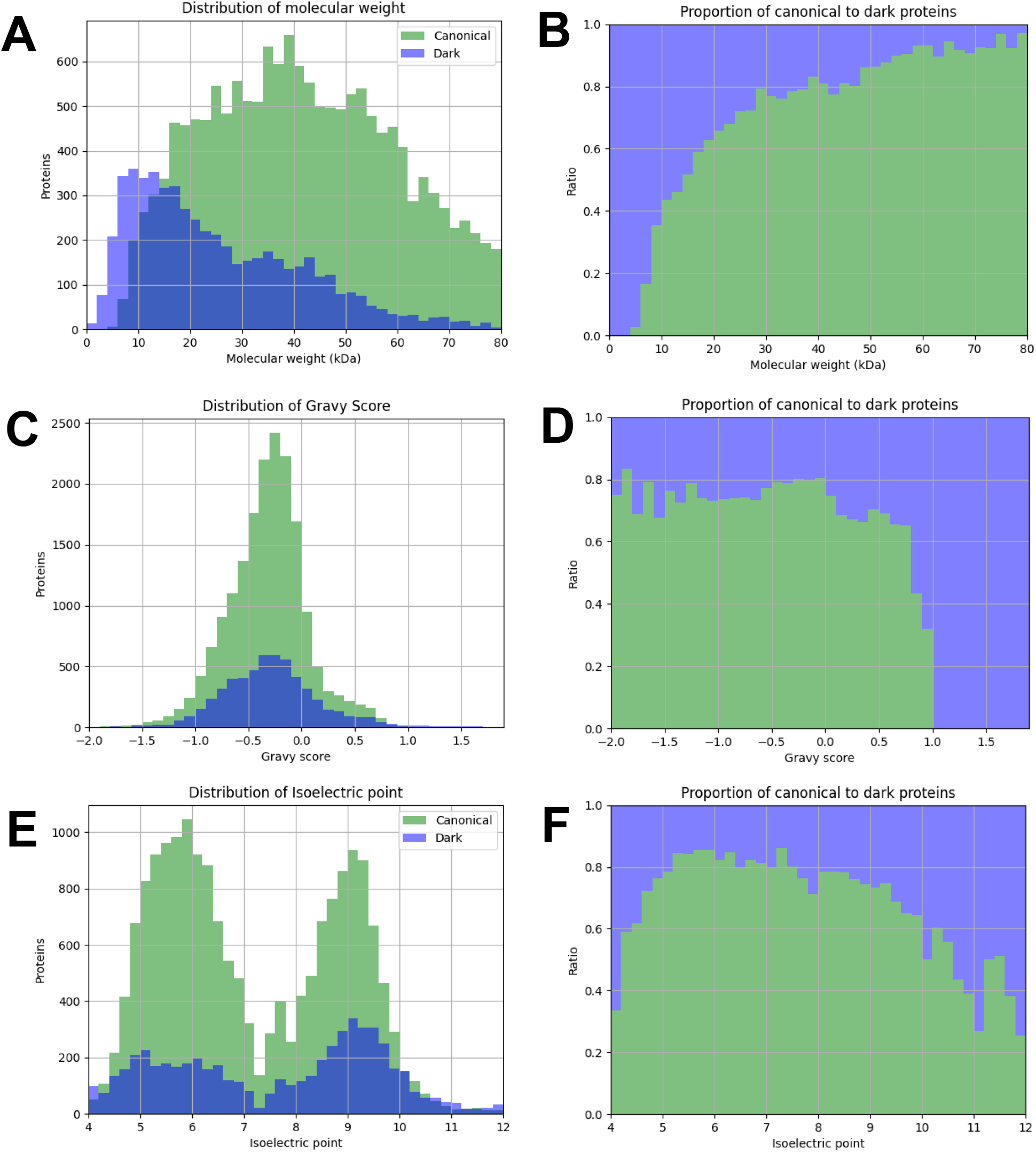
Distributions of physicochemical properties of the 18079 canonical (green) and 5595 dark (purple) proteins. A,C,E, Absolute counts of proteins within each bin for canonical and dark proteins. B,D,F, The proportion of canonical and dark proteins within each bin. A,B. Distributions and proportions of the molecular weight (kDa) of canonical (green) and dark (purple) proteins. Proteins with molecular weights between 0 and 80 kDa are shown. C,D. Distributions and proportions of the hydrophobicity (gravy score) of canonical (green) and dark (purple) proteins. Proteins with gravy score between -2.0 (hydrophilic) and 2.0 (very hydrophobic) are shown. E,F. Distributions and proportions of the isoelectric point (pI) of canonical (green) and dark (purple) proteins. Proteins with pI between 4.0 (acidic) and 12 (very basic) are shown.

In addition to these inherent properties of the canonical and dark proteins, we also explored the distributions of computed RNA abundances of the transcripts across 5,673 single and paired-end RNA-seq quality-controlled and filtered datasets from (Palos et al., 2022) with reads aligned to the Arabidopsis genome (see Materials and Methods). We excluded 345 protein coding genes that were never expressed above the median, as well as 309 undetected genes from the remaining analyses which were likely undetected due to mapping limitations with overlapping or highly similar genes (Supplemental Data Set S2). To evaluate mRNA expression patterns for the canonical and dark proteins, we considered two metrics, *i.e.* the percentage of RNA-seq data sets in which the transcript for a gene was detected (Figure 5A,B) and the maximum transcripts per million (TPM) for each expressed gene in any one of the RNA-seq data sets (Figure 5C,D). Figure 5A displays the distribution of the percentage of the 5673 RNA-seq datasets in which each transcript was detected. The highest bin (99-100%) is truncated at 1000 genes to better show details of the other bins (the true height of this highest bin is 12000). The relative proportions of canonical and dark proteins in each transcript bin are more easily seen in the proportion plot (Figure 5B) which shows that the proportion increases linearly across most of the range of transcript detection, except for the extremes at the ends. In other words, the more often a transcript for a gene is detected in one of the RNA-seq datasets, the higher the chance that the protein is canonical. For genes where this RNA detection percentage was below ∼5%, the predicted protein was typically not detected (*i.e*. dark), whereas for genes where the transcript was detected in >98% of the RNA-seq datasets, the predicted protein was nearly always canonical. Figure 5C depicts the distribution of the highest TPM among the analyzed RNA-seq experiments for each of the canonical and dark proteins. The TPM values extend as high as 207,000 (for seed storage protein albumin 3 - At4G27160) but the proportion does not change substantially above 100 TPM, and we only depict the range 0 to 500 TPM. Clearly the proportion of dark proteins rapidly increases when the maximum TPM falls below ∼100 TPM, suggesting that transcript abundance is likely influencing the detectability of proteins in MS analyses.

**Figure 5.**
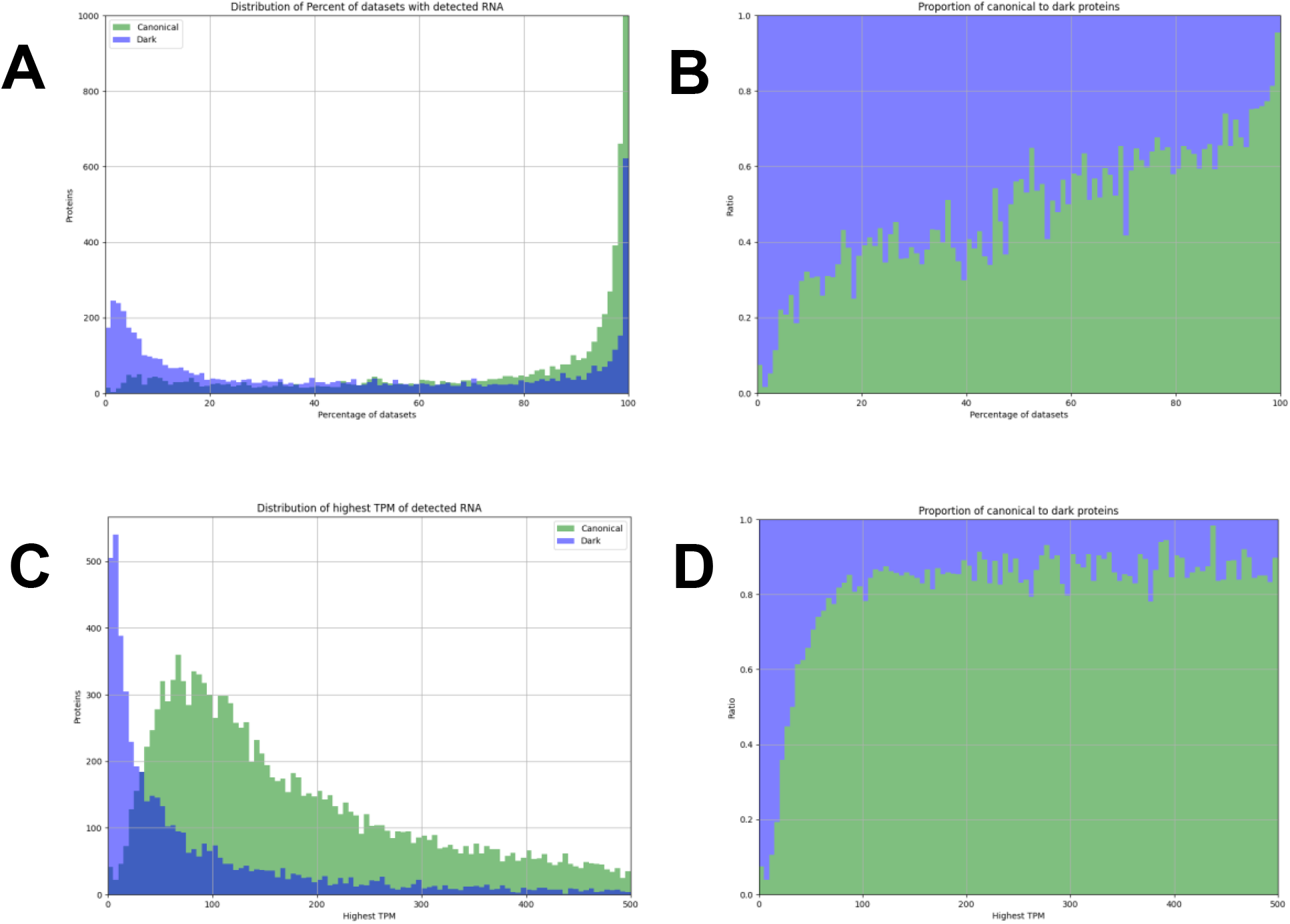
Transcript abundance and observation frequency of 26975 nuclear-encoded protein coding genes in 5673 high quality RNA-seq datasets. A,B, Distributions of the percentage of RNA-seq datasets with detected transcripts associated with the canonical (green) and dark (purple) proteins. A, Absolute counts of proteins within each bin and B, proportion of light and dark proteins within each bin. C,D, Distributions of the maximum transcripts per million (TPM) among all RNA-seq experiments for the detected transcripts associated with the canonical (green) and dark (purple) proteins. Absolute counts of proteins within each bin (C) and the proportion of light and dark proteins within each bin (D). The number of TPM extends as high as 207,000 for seed storage protein albumin 3 (AT4G27160), followed by seed storage cruciferin 1 and 3 (AT5G44120 and AT4G28520), Rubisco small subunit 1A (AT1G67090) and the hypothetical very small (33 aa) protein AT2G01021.

### Machine learning models to predict and understand MS-based detection of Arabidopsis proteins

Figures 4 and 5 showed that each of the protein and RNA attributes has a substantial influence on whether proteins are canonical or dark. Taking advantage of these attributes to better understand why these dark proteins are not observed, we trained both an artificial neural network (ANN) model and a random decision forest (RDF) model for the canonical and dark proteins based on physicochemical protein properties and RNA expression patterns. The quantitative output of these models was the probability for proteins to be canonical. The starting point was a table of 18079 nuclear-encoded canonical proteins and 5595 nuclear-encoded dark proteins for a total of 23674 proteins (uncertain and redundant proteins are left out for the training of the models; proteins without RNA values are also left out), as well as the computed physicochemical and RNA attributes discussed above. Figure 6 shows the receiver operating characteristic (ROC) curves to visualize the RDF (A,B) and the ANN (C,D) model performances trained on each of the features individually and collectively. ROC curves measure the ability of the model to distinguish between canonical and dark proteins. Figure 6E shows that the % detected transcript made the most important contribution to the RDF model followed by highest TPM and molecular weight. The overall accuracy of the RDF model when trained on all attributes was slightly better to the ANN model with area under the curve (AUC) values of 0.94 vs 0.91. Both the RDF and ANN models were robust as their ROC curves did not depend on which subset of the input data was used for training (Figure 6C,D). Supplemental Data Set S9 provides the protein and RNA features (input) for the models as well as the output (probability to be canonical).

**Figure 6.**
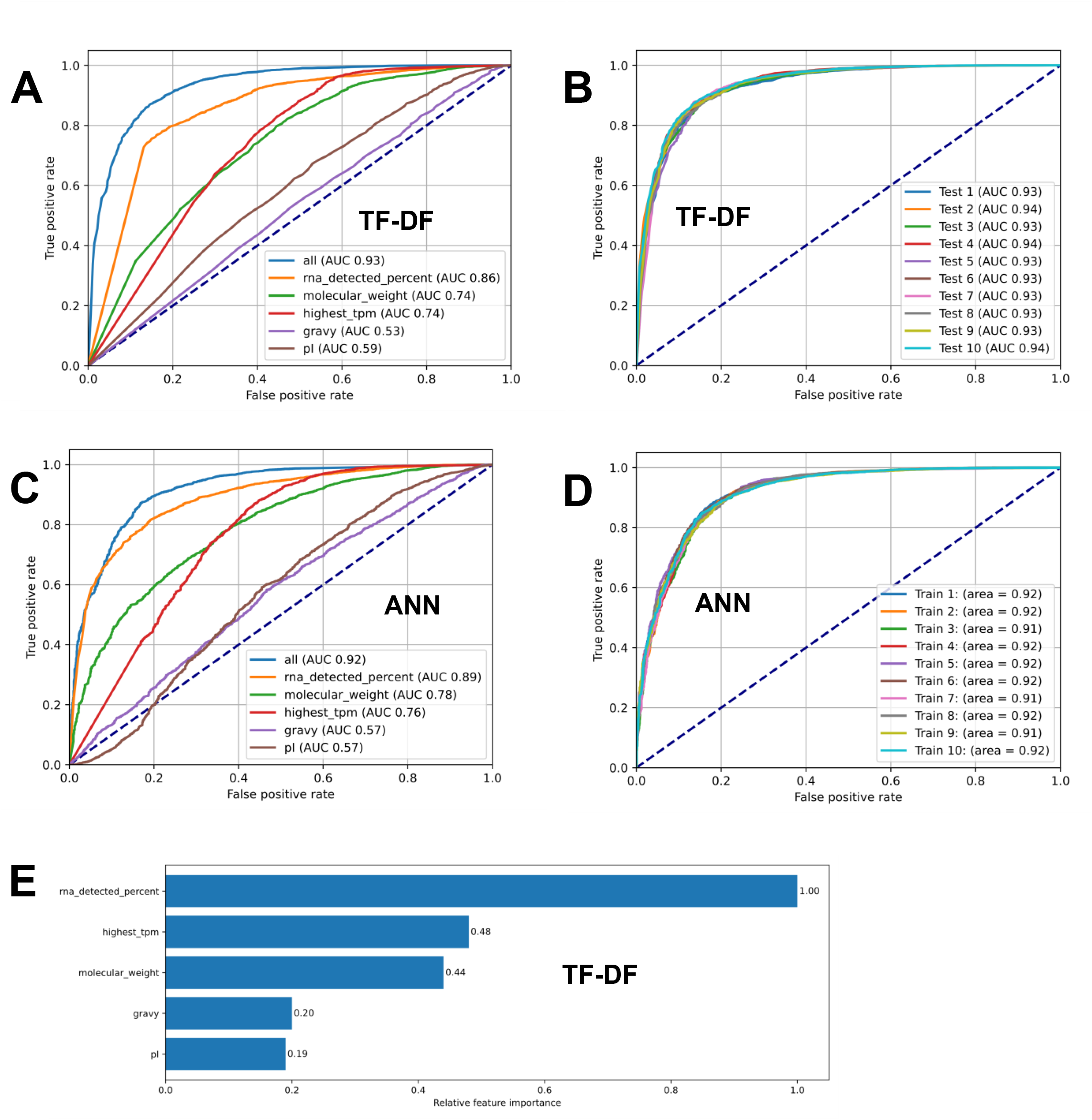
Machine learning models (ANN and TF-DF) to predict the probability of Arabidopsis proteins to be detected at the canonical levels in build 2. A-D, ROC curves for TF-DF models (A,B) or ANN (C,D) models trained on protein physicochemical properties and RNA expression data. A higher percentage of area under the curve (AUC) signifies better accuracy whereas an AUC of 0.5 (denoted by the dotted navy line) signifies near random prediction. As shown, % RNA detected, molecular_weight, and highest TPM enhance the performance of an ANN model, whereas pI and gravy barely impact it. B,D, ROC curves of TF-DF (B) and ANN (D) models trained on 10 randomized subsets of the same size from the input data. The accuracy of the TF-DF and ANN models are consistently around 93% and 92%, respectively. E, Feature importance. The TF-DF model has several built-in methods that calculate the significance of features to a model’s performance.

Even though the AUCs in the ROC curves were high, there is a substantial number of predicted canonical proteins that were in fact dark proteins and vice versa. To better understand possible reasons for these false predictions (outliers), we assembled two sets of outliers using the combined outcomes of both machine learning models, as follows: For dark protein outliers (predicted to be canonical, but dark), we required that both models calculated a probability (to be canonical) of >0.80; this resulting in 222 outliers. These outlier dark proteins had average physiochemical properties (47 kDa, –0.4 Gravy, 7.3 pI) and moderate average RNA expression values (96% RNA detected, highest TPM 361). Hence these undetected proteins appeared to have favorable properties (not very low molecular weight, not hydrophobic, not very basic and significant transcript levels and detection across RNA-seq datasets), yet were not detected by MS. For canonical protein outliers (predicted to be dark, but canonical) we required that both models calculated a probability (to be canonical) of <0.20; this resulted in only 42 outliers; these outliers had the average physiochemical properties of 24 kDa MW, –0.3 Gravy, 7.9 pI and low average RNA expression values (19% RNA detected, highest TPM 33). Hence these unexpected canonical proteins have very low transcript levels and were often not detected in RNA-seq experiments yet were detected at high confidence levels. We then further explore the underlying scenarios for this unexpected behavior based on functional annotations and manual inspection, as described in a section further below (*Explanations for unexpected canonical or dark proteins*).

### Biological properties and functions of the dark proteome

Based on the description of the proteins in TAIR, we observed that proteins annotated as ‘hypothetical proteins’ (some have DUF domains) were highly overrepresented at 24% of all dark proteins (1349 out of 5595), compared to just 2.6 % of the canonical proteins (476 out of 18079) (Figure 7A). These hypothetical proteins are annotated in TAIR as ‘protein coding’ and not as pseudogenes. On average, the predicted observability to be canonical for these hypothetical proteins was indeed much lower for the dark proteins than the canonical proteins (Figure 7B). Proteins annotated as ‘unknown’ and/or proteins with a DUF domain’ represented 5% of the dark proteins and 4.3% of the canonical proteins (Fig. 6A), thus lacking this overrepresentation in dark proteins.

**Figure 7.**
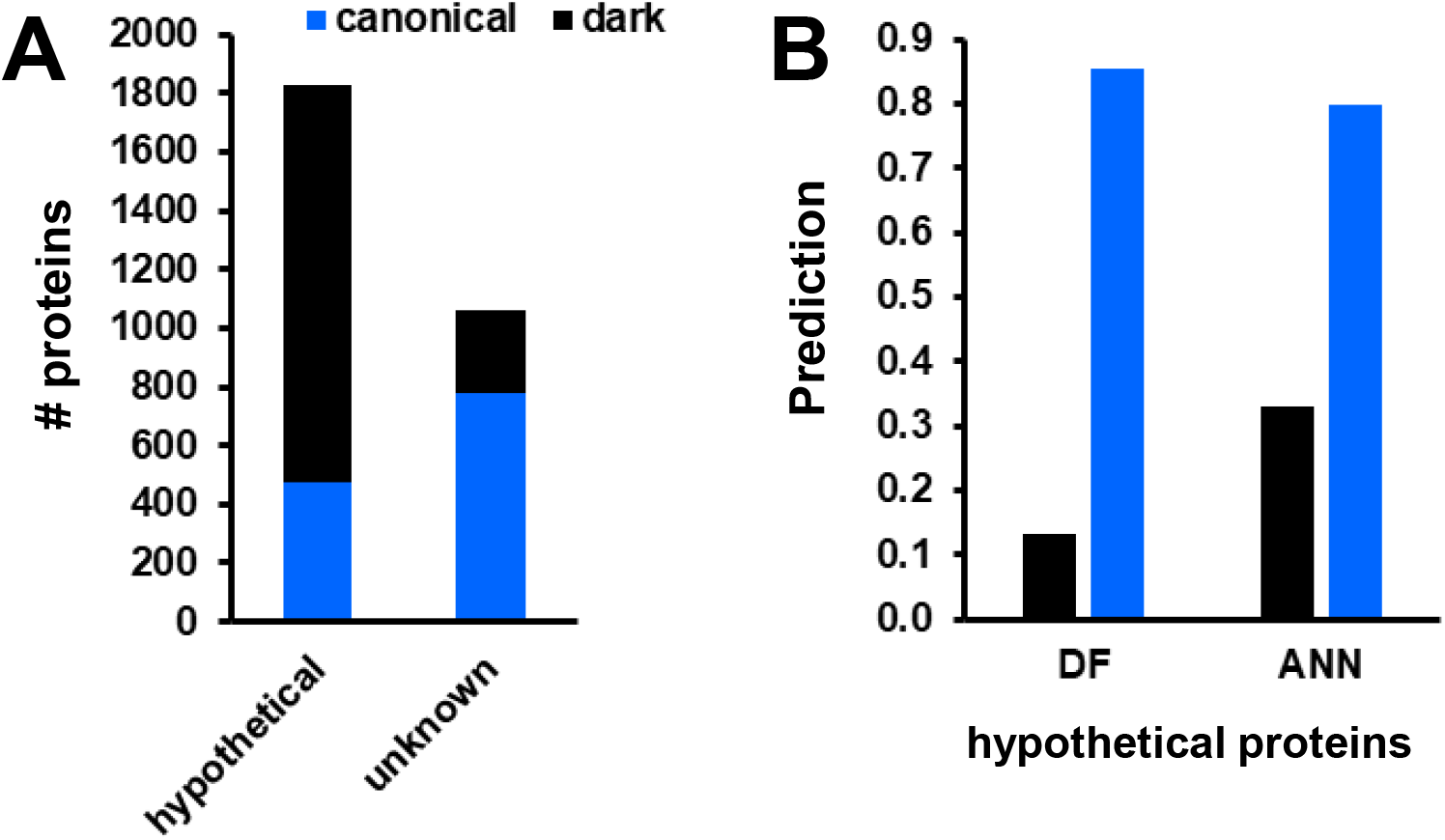
Hypothetical and unknown/DUF proteins in the dark and canonical proteome and their predictions to be canonical. All canonical and unobserved proteins were scored for the presence of the words “hypothetical”, “unknown” or “Domain of Unknown Function (DUF)” in their description from Araport11/TAIR. A, Hypothetical and unknown proteins in the dark and canonical proteome. B, Predicted observability for the hypothetical proteins to be canonical using the two machine leaning models (DF and ANN).

To take an unbiased approach to determine if the dark proteome is enriched for particular types of proteins, we used the Arabidopsis Gene Ontology (GO) enrichment analysis (Ashburner et al., 2000; Ge et al., 2020; Gene Ontology, 2021) for the three GO categories Biological Process (BP), molecular function (MF) and cellular component (CC). GO analysis was done by comparing all dark proteins to either the sum of canonical and dark proteins or all predicted Araport11 proteins; the results were similar for both comparisons, and we show therefore the results of the latter. We did not observe any significant enrichment for the CC categories suggesting that the build #2 did not under-sample any particular subcellular localization. Indeed, the PXDs that are included in build #2 deliberately include all plants parts, and most subcellular fractions such as chloroplasts, mitochondria, etc. However, significant enrichment was observed for BP and MF with the 20 most significant GO terms (lowest FDR) for BP or MF shown in figure 7A,B. A protein can have several GO terms for each category and different GO terms can relate to similar processes or functions (Supplemental Data Set S10). There were 520 proteins in the top20 GO terms for BP and 739 proteins for the top20 GO terms for MF, with 271 found in both.

Upon analysis of the enriched BP GO terms (Figure 8A) and the protein IDs, we determined that there are mainly three broad types of protein functions enriched in the dark proteome. These are: i) <u>149 signaling peptides/peptide hormone</u>s such as members of the clavata family, defensins, root meristem growth factor (GO terms: Cell signaling (involved in cell fate commitment), Cell-Cell signaling, Cell fate commitment, Signaling receptor activity, Signaling receptor binding, Regulation of asymmetric cell division, nitrate import, cell killing, killing of other cells of other organisms, phloem development, regulation of cell differentiation), ii) ∼<u>236 proteins involved in the ubiquitination pathway</u>, including 160 E3 ligases, one E2 conjugating enzyme, 8 ubiquitin(-like) proteins (Go terms: Protein ubiquitination, protein modification by small conjugation (or removal), Ubiquitin(-like) protein ligase activity, Positive regulation of (proteasome) ubiquitin-dependent protein catabolic process), iii) <u>∼130 proteins associated with DNA & RNA related processes</u> (GO terms: RNA/Nucleic acid phosphodiester bond hydrolysis (endonucleolytic), RNA-dependent DNA biosynthetic process, DNA biosynthetic process). Many of these proteins belong to superfamilies such as: RNA-directed DNA polymerase (reverse transcriptase)-related family (it is not clear what function these have in Arabidopsis), non-LTR retroelement reverse transcriptase, reverse transcriptase zinc-binding protein, Polynucleotidyl transferase ribonuclease H-like superfamily, ribonuclease H superfamily polynucleotidyl transferase. Many of these proteins seem to have no defined function.

**Figure 8.**
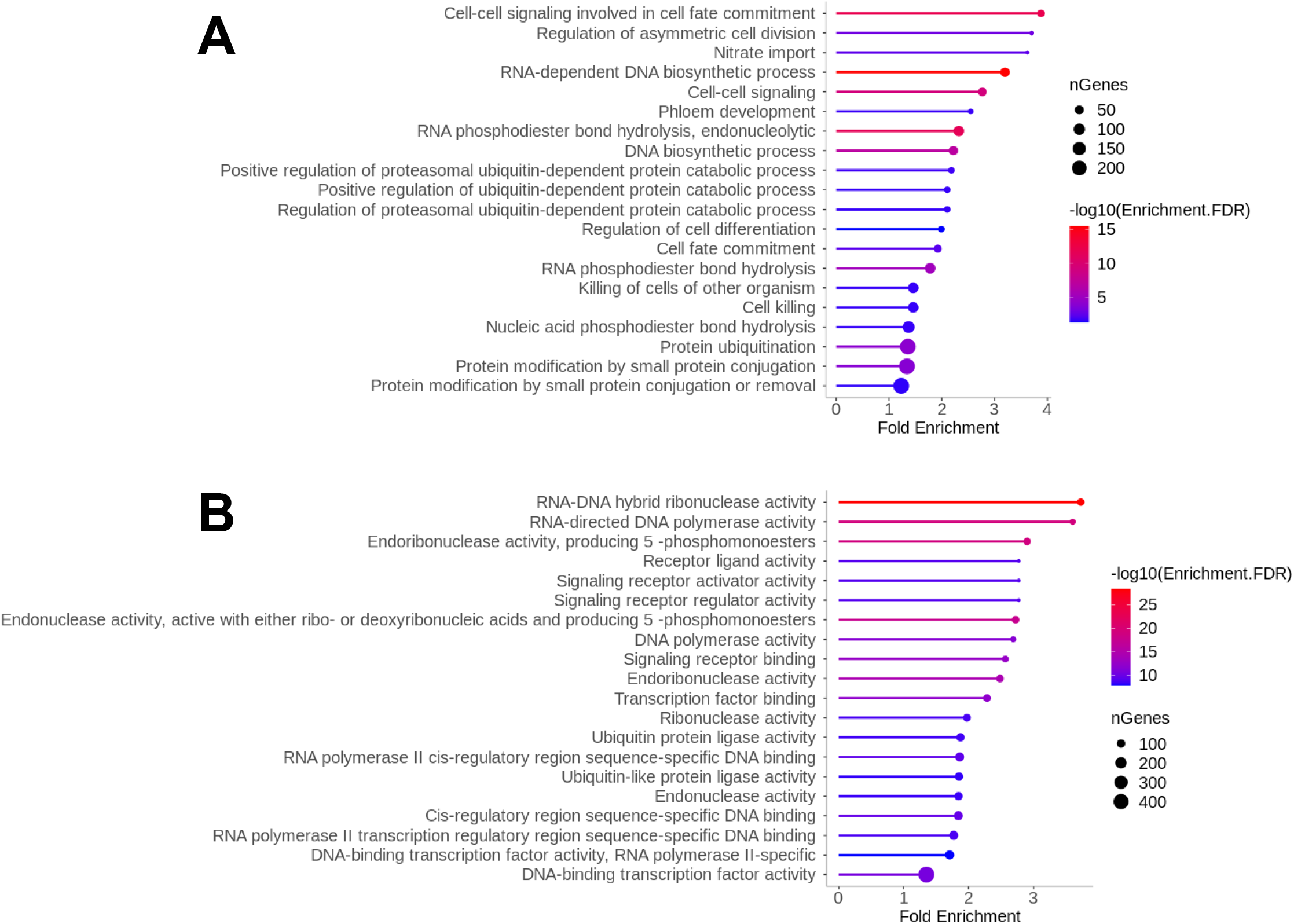
GO enrichment of the 5595 dark proteins compare to all predicted Arabidopsis proteins for Biological Process and Molecular function. A,B, The 20 most significant GO terms (lowest FDR) are shown, ordered by fold enrichment for biological process (A) and molecular function (B)

Analysis of the top 20 enriched MF GO terms (Figure 8B) showed 115 UBI-related proteins and 70 signaling peptides as in the BP GO analysis above. But transcription factor proteins represent by far the most enriched molecular function, with a total of over 400 members of different TF families (*e.g.* AP2/EREBP, ARF, Auxx/IAA, bHLH, bZIP, C2C2(Zn), C2H2, MADS box, MYB, CCAAT, WRKY) (GO terms: DNA-binding transcription factor activity, Transcription factor binding, RNA polymerase II cis-regulatory region sequence-specific DNA binding, Cis-regulatory region sequence-specific DNA binding, RNA polymerase II transcription regulatory region sequence-specific DNA binding, DNA-binding transcription factor activity, RNA polymerase II-specific, DNA-binding transcription factor activity). The 2^nd^ largest molecular function was for various endonuclease activities with ∼83 proteins, including several types of reverse transcriptases and ribonuclease H family members (GO terms: Endonuclease activity, Endonuclease activity, active with either ribo-or deoxyribonucleic acids and producing 5 - phosphomonoesters, Ribonuclease activity, Endonuclease activity, RNA-DNA hybrid ribonuclease activity). Finally, there were 27 proteins associated with the GO terms RNA-directed DNA polymerase activity and DNA polymerase activity; most of these were also annotated as reverse transcriptases.

### Signaling peptides/peptide hormones are highly over-represented in the dark proteome

The GO enrichment analysis (Fig. 8A,B) suggested that proteins encoding for plant signaling peptides and/or peptide hormones are strongly overrepresented in the dark proteome. Most are inactive precursors (preproproteins of ∼7 to ∼12 kDa) that undergo a multistep proteolytic processing to result in the relatively small (between ∼5 and ∼100 amino acids) bioactive peptide signals (Matsubayashi, 2014; Tavormina et al., 2015; Olsson et al., 2019; Stintzi and Schaller, 2022). These small proteins are of great importance in many aspects of plant life. Most of these precursors are secreted through a cleavable N-terminal signal peptide (sP) for targeting into the ER, followed by traveling through the Golgi, plasma membrane and into the apoplast. However, the mode of bioactive peptides can be extracellular or intracellular. We note that there are also bioactive peptides derived from different types of short open reading frames (sORFs, uORFs, lncRNA, pri-miRNA), most of which do not yet have an ATG identifier in the current TAIR annotation (Takahashi et al., 2019; Hu et al., 2021). Bioactive plant peptides have traditionally been grouped into (i) cysteine-rich peptides that form internal disulfide bonds, and (ii) post-translationally modified small peptides that undergo one or more PTMs during their passage through the ER or Golgi (*e.g.* tyrosine sulfation (Kaufmann and Sauter, 2019), proline hydroxylation, *etc*) (Matsubayashi, 2014; Olsson et al., 2019).

Many peptide families have been recognized (Matsubayashi, 2014; Olsson et al., 2019; Kim et al., 2021; Stintzi and Schaller, 2022), including Clavata/embryo-surrounding region (CLE) (Willoughby and Nimchuk, 2021; Yuan and Wang, 2021), Epidermal Patterning Factor (EPF) (Yuan and Wang, 2021), phytosulfokine-alpha (PSK) (Matsubayashi, 2014), cysteine-rich peptides of the LURE family (Zhong et al., 2019), Embryo Surrounding Factor (ESF), PAMP-induced secreted peptides (PIP), Plant Peptides containing Tyrosine sulfation family (PSY) (Tost et al., 2021), root meristem growth factor (RGF), caesarian strip integrity factor (CIF) (Fujita, 2021), inflorescence deficient in abscission (IDA), precursor of plant elicitor peptide (PROPEP) (Huffaker et al., 2006; Bartels et al., 2013), defensin-like (DFL) and POLARIS which is not part of a larger family. We assembled a tentative list of their protein ATG identifiers (330 genes) to get a better understanding to what extent they were identified in the new PeptideAtlas build (Supplemental Data Set S11). PeptideAtlas identified 92 (28%) at the canonical level and 144 (44%) were part of the dark proteome (Figure 10A). The remainder of these 330 proteins were identified at various lower confidence levels often as part of a group of homologs (48 weak, 2 insufficient evidence, 14 subsumed, 14 marginally distinguished, 6 indistinguishable representative) (Figure 9A). The identification level within each family (Fig. 9B,C) shows that the majority of members of some families were identified at the canonical level (PEP, CAP, LTP and THIONIN), whereas the identification rate in other families was very low (CIF, CLE, CEP, EPF, IDA, PAMP, PSY, RTFL/DVL, RGF) with >64% members unobserved (dark). The correlation between average (or median) precursor length for each family and identification status is weak. This is logical because these proteins are synthesized as precursors followed by one or more proteolytical cleavages. Furthermore, for family members decorated with PTMs on the amino acid residues Y, S or P (see Figure 9B) identification rates should be lower since our database search does not include these PTMs because they are relatively rare. Inclusion of such PTMs in regular searches is not appropriate and would result in many false discoveries.

**Figure 9.**
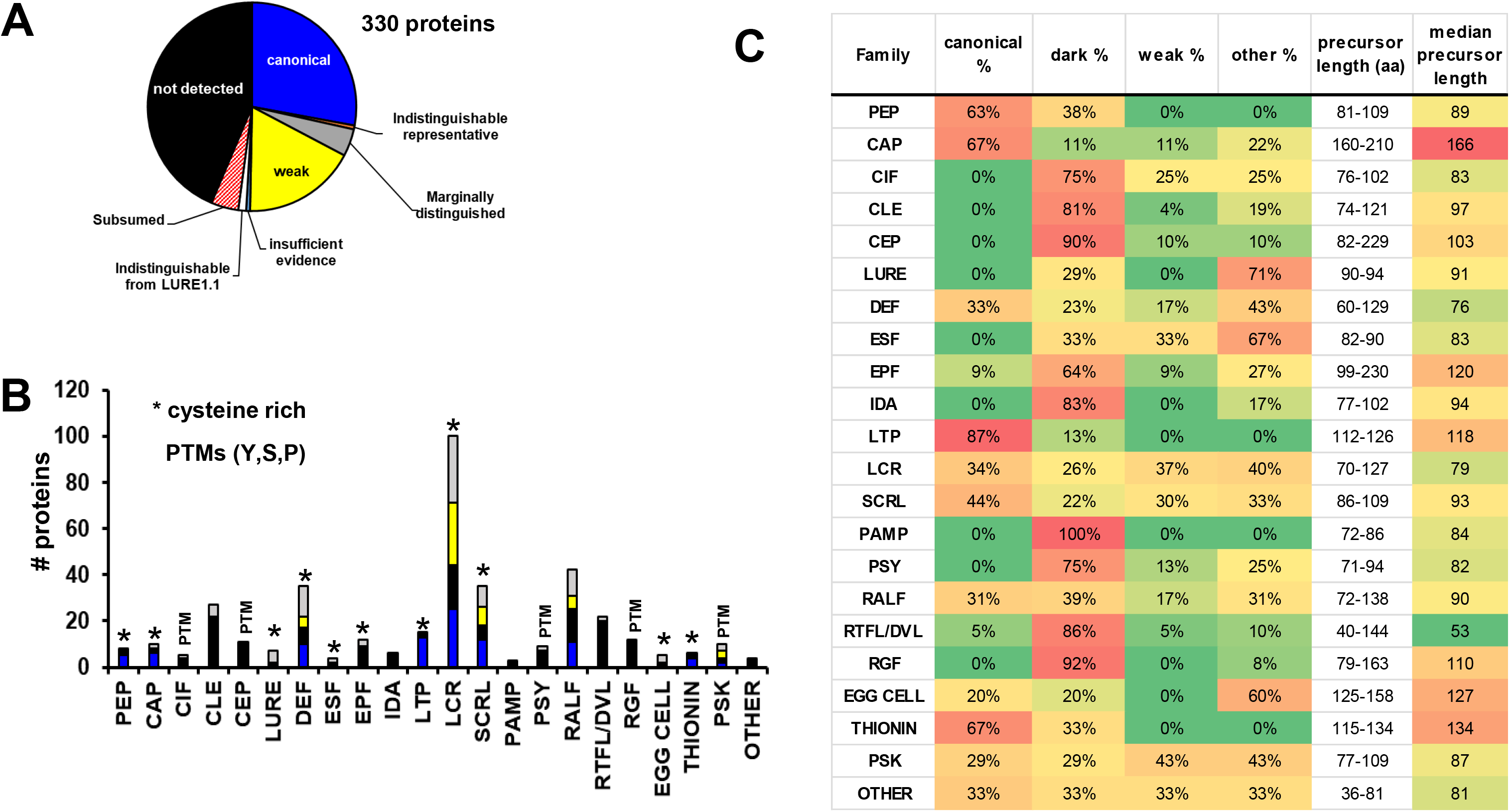
Identification status of members of different signaling peptide families in build 2. A, Overall identification status across 8 confidence tiers of the 330 signaling peptide producing proteins (Supplemental Data Set S11). The tiers system is described in more detail in (van Wijk et al., 2021). Identified protein with status ‘weak’ have at least one uniquely mapping peptide of 9 amino acid residues but does not meet the criteria for canonical (at least 2 uniquely mapping non-nested peptides of at least 9 residues with at least 18 residues of total coverage). B, Bar diagrams of proteins within each of the peptide signaling families. Color coding within each bar indicates the number of proteins not-observed (black), weak (yellow), canonical (blue) or in other tiers (gray). * indicates cysteine rich peptides. PTMs indicates known presence of PTMs of signaling peptides. C, Listing all families, identification level and precursor length (range and median)

**Figure 10.**
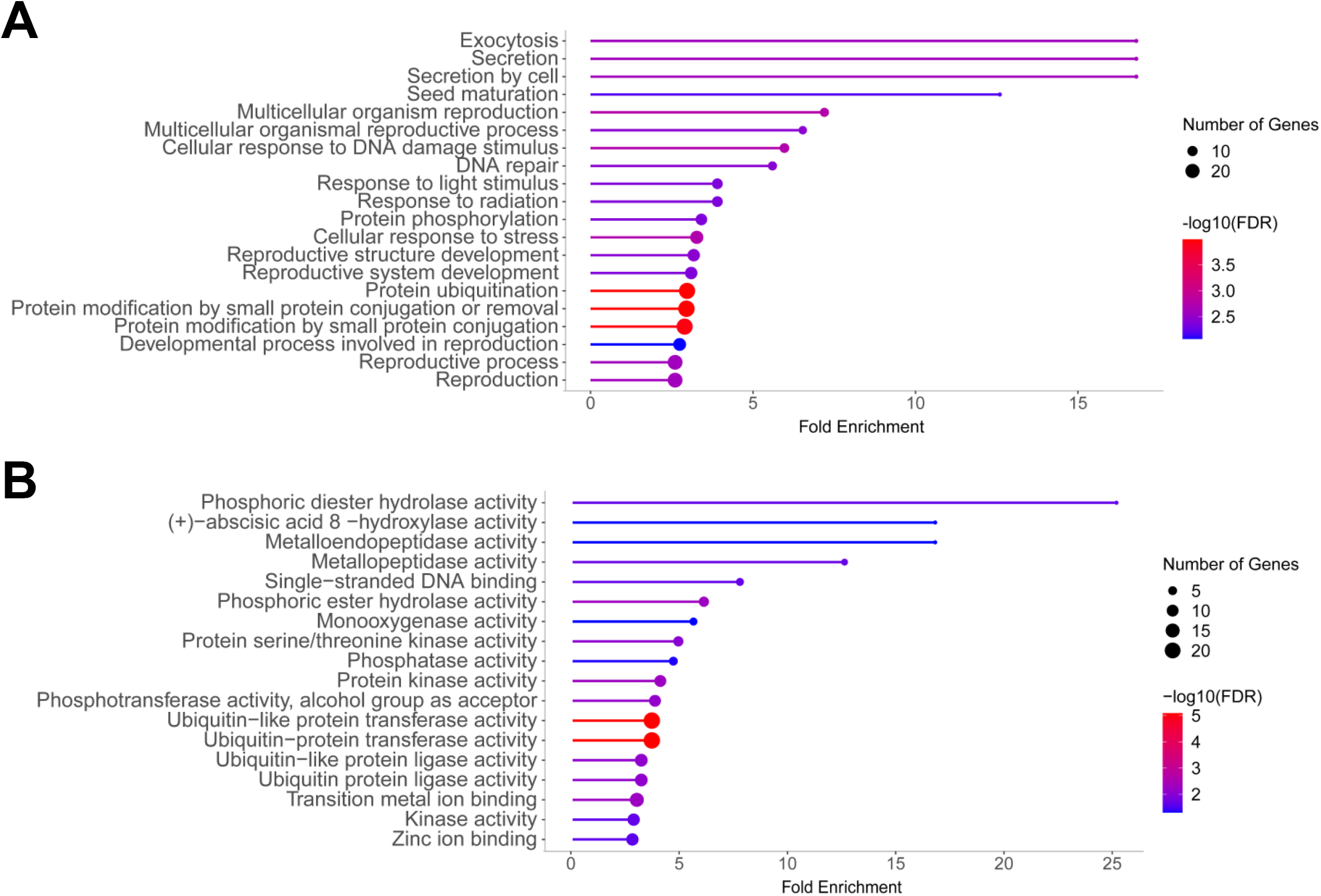
GO enrichment of 222 outlier dark proteins compare to all 5595 dark proteins or Biological Process and Molecular function. The outliers are defined as dark proteins having a predicted probability to be canonical of >0.8 by both machine learning models. A,B, The 20 most significant GO terms (lowest FDR) are shown, ordered by fold enrichment for biological process (A) and molecular function (B).

Interestingly, PSMs of the identified proteins ranged from just a few to several thousand for several LTP family members (LPT1,2,3,4) and DEF members (PDF1.1, 1.2A/B/C. 1.3). Sequence coverage was > 60% for some 22 preproteins, including several THIONINS, CAPs and a few PEPs; further close inspection of the matched peptides in PeptideAtlas showed that the sequence coverage started downstream of the cleavable signal peptide and mostly or completely included the predicted C-termini. More biological insight into the accumulation and maturation of these signaling peptides can be derived by exploring the associated metadata (stored and linked in PeptideAtlas) and relate that to identification status, protein coverage and abundance as measured by PSMs in PeptideAtlas. Identifications of the unobserved and low confidence peptides will require targeted experimental approaches, and specific search strategies (*e.g.* allowing for specific PTMs).

### E3 ligases are highly over-represented in the dark proteome

The GO enrichment, and inspection of the associated protein IDs, showed that E3 ligases were over-represented in the dark proteome. Arabidopsis has some ∼1400 E3 ligases that each target one or several substrates for polyubiquitination and subsequent degradation by the proteasome. The required amount of an E3 ligase in a cell greatly depends on the number and abundance of its substrates. The dark proteome included 601 E3 ligases (10.7% of the dark proteome) whereas the canonical proteome included 429 E3 ligases (2.4% of the canonicals). Comparing the dark and canonical E3 ligases shows that these 2 groups do not differ in the three physicochemical properties (size, gravy, pI) but that dark proteins have on average much lower transcript levels (both TPM and % observed).

### Proteins with short half-life or extensive proteolytic processing - protein features not considered in the machine learning

There are two protein features (attributes) that were not considered in the machine learning models. These features are i) proteolytic trimming of the preproteins or (pre)proproteins, and ii) short protein half-life resulting in net low abundance under most conditions. Both scenarios make it harder to detect such proteins by MSMS than predicted by the machine learning models. We already described examples for extensive proteolytic trimming for plant signaling peptides/peptide hormones which are indeed overrepresented in the dark proteome.

Proteins that are predicted to be canonical but with a conditional short-half life might go undetected (dark proteins) or with very low number of PSMs, because they are continuously degraded under most circumstances. However, the half-life of most proteins is unknown. One of the exceptions is the set of five transcription factors in the group VII of the Ethylene Response Factor (ERF-VII) family involved in oxygen sensing (Gibbs et al., 2015; van Dongen and Licausi, 2015; Hammarlund et al., 2020; Weits et al., 2021; Barreto et al., 2022) (Table 6). These proteins have a short half-life under normal oxygen concentration (normoxia) because they are degraded by the proteasome through the N-degron pathway but become stabilized during hypoxia or anoxia. These proteins have a cysteine in the 2^nd^ position from the N-terminus. After removal of the start methionine by methionine amino peptidases, these N-terminal cysteines are enzymatically oxidized by APs by Plant Cysteine Oxidases (PCDs) which is then followed by enzymatic arginylation (*i.e.* additional of an arginine residue) (White et al., 2017; Hammarlund et al., 2020). The arginylated N-terminus is then recognized by specific E3 ligases, resulting in polyubiquitination and degradation by the proteasome. At low cellular oxygen concentrations (hypoxia) due to respiration or environmental conditions (*e.g.* flooding, high altitude), these transcription factors stabilize because the enzymatic oxidation is slowed down (Abbas et al., 2022). In Arabidopsis there are five members of this ERF-VII family, *i.e.,* hypoxia response 1 (HRE1; AT1G72360), HRE2 (AT2G47520), related to apetala 2.12 (RAP2.12; AT1G53910), RAP2.2 (AT3G14230), RAP2.3 (AT3G16770).

**Table 6.** PeptideAtlas detection of the ERFVII transcription factor members involved in oxygen sensing.

|  |  |  |  |  |  |  |  |  |  |  |  |
| --- | --- | --- | --- | --- | --- | --- | --- | --- | --- | --- | --- |
| Accession | name | PA Status | # PSMs & plant materials | MW | PI | GRA VY | % rna detecte d | average TPM | highest TPM | probabilit y DF | probability ANN |
| AT3G16770.1 | RAP2.3 | Weak | one Phosphopeptide – 2 PSMs (cell culture-phospho, callus-phospho) | 27.76 | 5.21 | -0.73 | 99.98 | 329 | 7877 | 0.997 | 0.939 |
| AT3G14230.1 | RAP2.2 | Marginally Distinguished AT1G53910.1 | two peptides – total 2 PSMs (cell culture) | 42.53 | 4.91 | -0.78 | 100.00 | 136 | 1932 | 1.000 | 0.969 |
| AT2G47520.1 | HRE2 | Weak | one peptide – 3 PSMs (cell culture, callus) | 19.35 | 6.41 | -0.86 | 82.78 | 7 | 1202 | 0.883 | 0.793 |
| AT1G72360.1 | HRE1 | Weak | one peptide – 2 PSMs (cell culture, flower) | 23.66 | 4.83 | -0.73 | 99.59 | 16 | 1375 | 0.737 | 0.928 |
| AT1G53910.1 | RAP2.12 | Canonical | five peptides – total 5 PSMs (Cell culture) | 39.8 | 5.19 | -0.74 | 100.00 | 148 | 1538 | 0.990 | 0.972 |

Table 6 summarizes the PeptideAtlas findings and protein attributes this ERF-VII family. Whereas there was MSMS support for all five proteins, the number of PSMs was very low (between 2 and 5). All but one peptide was from callus or cell culture – callus is known to have low internal [O_2_] (Hammarlund et al., 2020) explaining why the proteins were observed in callus. It seems quite plausible that plant cell cultures also might experience hypoxia (due to high respiration and low/no photosynthesis). The ERVII TF proteins are predicted to be canonical (predicted observability between 0.7 and 1) (Table 6). However, only RAP2.12 was identified at the canonical level but only in one specific experiment using cell cultures (PXD013868, experiment 8213 https://db.systemsbiology.net/sbeams/cgi/PeptideAtlas/ManageTable.cgi?TABLE_NAME=AT_SAMPLE&sample_id=8213). Furthermore, RAP2.3 was only identified with a phosphorylated peptide identified in callus and in cell cultures. Their transcripts were detected in the majority (> 82%) of the 5673 RNA-seq datasets and all proteins have very high maximum TPM values (1202-7877). This is a nice example where the correlation between predicted probability to be canonical (from the machine learning models) and observed overall number of PSMs suggest unusual properties of the proteins, in this case short half-life. The associated metadata help to provide biological context as the findings for these ERF-VII proteins illustrate.

### Explanations for unexpected canonical or dark proteins

A small subset of dark proteins (222 out of 5595) were predicted by both machine learning models to be canonical (p>0.8) and 44 canonical proteins were predicted to be dark (p<0.2). To explore biological scenarios for these unexpected dark or canonical proteins we used both GO enrichment and manual evaluation. We compared GO distributions of the 222 dark outliers and the 5595 dark proteins (Figure 10 and Supplemental Data Set S10). The highest number of proteins were found for GO terms associated with <u>ubiquitination</u> (Protein ubiquitination, protein modification by small conjugation (or removal), Ubiquitin(-like) protein transferase activity, Ubiquitin(-like) protein ligase activity). Upon further inspection these were mostly E3 ligases, in particular RING ligases. Other GO terms pointed to enrichment in kinases, terms associated with reproduction, DNA repair, and response to light stimulus or response to radiation, but the genes associated with these GO terms have quite broad range of functions (*e.g.* transcription factors, some E3 ligases).

Because the number of unexpected dark proteins was relatively small, we explored these also manually. Two of the unexpected dark proteins were chloroplast sigma factors 1 and 3 (SIG1 and SIG3; AT1G64860 and AT3G53920) with predicted probability to be canonical between 0.84 and 0.98. Both are very basic proteins (9.5 and 9.8 pI) with have relatively high molecular weight of the precursors (56 and 65 kDa) and were detected in nearly all 5673 RNA-seq data sets with the highest TPM of 383 and 105; hence it is therefore surprising that they were not detected by MSMS. Arabidopsis has six sigma factors (SIG1-6) (Chi et al., 2015; Puthiyaveetil et al., 2021) and also SIG4 and SIG5 were unobserved (but with lower probabilities to be canonical than the other sigma factors), whereas SIG2 and SIG6 were canonical. Protein sequence coverage by matched peptides for SIG2 and SIG6 were 45% and 20%, respectively with 16 and 7 PSMs respectively, showing that also SIG2 and SIG6 are of low general abundance. The most logical explanation is that the half-life of all sigma factors is relatively short. Chloroplast GUN1 (AT2G31400) is a large PPR protein (100 kDa) is known to have a short half-life of just several minutes because it is degraded by the Clp chaperone-protease system (Wu and Bock, 2021). GUN1 was identified at the canonical level with 12% sequence coverage but only 9 PSMs which is relatively low given its large size and high TPM (596). These examples serve to show dark proteins with a predicted probability to be canonical are likely enriched for protein with short-half-life or have unique expression patterns.

### Lessons from new PXDs in build 2 that contribute most effectively to identifying new canonical proteins

To inform possible strategies to efficiently identify the remaining 21% of the predicted proteome, we evaluated which of the new PXDs that we had selected to generate the new build had the most impact. Figure 11 shows the relation between the number of identified spectra and newly identified canonical proteins (not identified at the canonical level based on earlier datasets) for each of the 63 new PXDs that we added for build 2. Six PXDs that each added the most new canonical proteins are annotated in the figure, together identifying 146 new canonical proteins. Reviewing these new proteins within each of these six PXDs for protein features, including function and molecular weight, identified clear patterns consistent with sample types.

**Figure 11.**
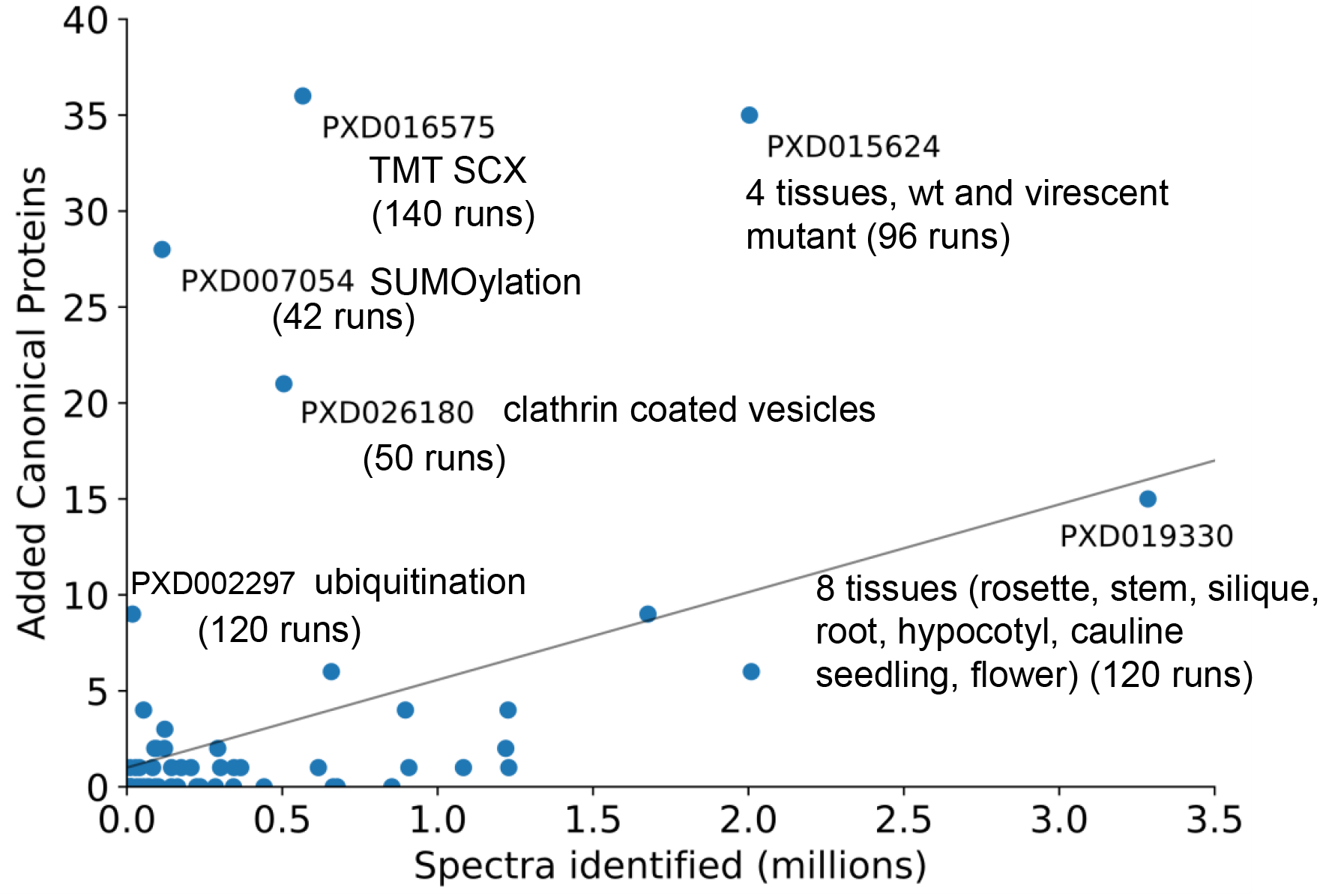
The relation between the number of identified spectra and newly identified canonical proteins for each of the 63 new PXDs that we added for build 2. Key information of the sample type is shown. Newly identified canonical proteins are proteins that were not yet identified as canonicals in build 1 or PXDs in build 2 with lower number. MS instruments used are: PXD016575 – Q Exactive HF-X; PXD007054 - LTQ Orbitrap Velos; PXD026180 - LTQ, Q Exactive HF, Q Exactive and LTQ FT Ultra; PXD015624 – Q Exactive, PXD0119330 - Orbitrap Velos Pro; PXD0002297 – Q Exactive.

PXD002297 contained 120 MS runs using a Q Exactive instrument from which we matched ∼18 thousand MSMS spectra yielding 9 new canonical proteins. This study used COFRADIC technology to map ubiquitination sites reporting 3,009 ubiquitination sites in 1,607 proteins (Walton et al., 2016). In PXD007054 we identified only 0.11 million MSMS spectra based on 42 MS runs, but yet this resulted in 28 new canonical proteins. This study was focused on identification of SUMOylated proteins using a three-step purification protocol based on 6His-tag and anti-SUMO1 antibodies from 8-day old Arabidopsis seedlings expressing a 6His-SUMO1(H89-R) transgene in wt and SUMO E3 ligase mutants *siz1-2* and *mms21-1* (Rytz et al., 2018). Interestingly, the new canonical proteins were highly enriched for transcription factors (17 out of these 28). PXD015624 provided 96 MS runs from which we matched 2 million MSMS spectra resulting in 35 new canonical proteins (Berger et al., 2020). The experiments involved label free proteomics of rosettes and roots from 8 weeks old plants and 2 weeks old seedlings of wild-type and *nfu2* plants (small and virescent) using a standard workflow (four replicates) involving protein separation by SDS-PAGE (4 slices per lane, tryptic digestion) and an Q Exactive Plus mass spectrometer. More than half of these new canonical proteins were larger proteins over 55 kDa, including five LRR kinases (98-106 kDa) and the glutamate receptor 2.3 (101 kDa). From PXD016575 we identified 0.57 million MSMS spectra and 36 new canonical proteins from 140 MS runs. The experiments involved the analysis of seedlings of wt and the autophagy-deficient mutant atg2-2 upon consecutive, temporary reprogramming inducing stimuli ABA and flg2 (Rodriguez et al., 2020). The proteomics workflow involved SDS extracted total seedling proteomes, TMT labeling followed by SCX chromatography and standard nanoLC-MSMS using a Q Exactive instrument. The new canonical proteins from this set included 19 proteins below 20 kDa, including several RALF signaling peptides; these small proteins are often missed in SDS-PAGE separated samples. PXD019330 was a truly large-scale proteomics study sampling multiple tissue types (roots, leaves, cauline leaves, stems, flowers, siliques/seeds, whole plant seedlings) at different developmental stages (Bassal et al., 2020). A standard workflow was used involving protein separation by SDS-PAGE (5 slices per lane, tryptic digestion) and an LTQ-Orbitrap Velos instrument and notably a long C18 column (50 cm) and long (9 hrs) elution with a total of 120 MS runs. We matched 3.29 million MSMS spectra resulting in 15 new canonical proteins. These new canonicals included several chloroplast membrane proteins (FAX4 and Lil1.2), a nitrate transporter and two very small metallothioneins. PXD026180 contained 50 MS runs from four different MS instruments (LTQ, Q Exactive HF, Q Exactive, LTQ FT Ultra) from which we mapped 0.5 million MSMS spectra and yielding 21 new canonical proteins. This study analyzed purified clathrin coated vesicles (CCVs) from undifferentiated Arabidopsis suspension cultured cells using both SDS-PAGE and in-solution digests, followed by nanoLC-MSMS on the extracted peptides (Dahhan et al., 2022). These six PXDs utilize a wide range of methods and plant materials, some high affinity enriched (SUMOylation, ubiquitination, CCV) and others including a range of different plant parts. As expected, several of the new canonical proteins are involved in vesicle transport.

As this snapshot of six PXDs illustrates, the proteomics-MS workflows showed a wide range of techniques (*e.g*. from SDS-PAGE with in-gel digests, to in-solution digest, TMT labeling and SXC chromatography) in all cases followed by reverse-phase nanoLC-MSMS but with different generations of MS instruments. Considering the total number of matched MSMS spectra, those PXDs that used affinity enrichment based on specific PTMs or isolation of highly specialized subcellular structures, clearly identified the most new canonical proteins when normalized to the number of matched spectra. This suggests that the identification of the remaining 21% of the predicted Arabidopsis proteome will be most effective when this will also include targeting specific subcellular structures and specific PTMs.

## CONCLUSIONS AND PERSPECTIVE

This second release of the Arabidopsis PeptideAtlas is based on ∼259 million searched raw MSMS spectra from 115 PXDs and includes 21017 protein identifications based on ∼70 million matched spectra (PSMs) and nearly 0.6 million distinct matched peptides. Compared to the first release (van Wijk et al., 2021) this represents an increase of 78% more PSMs, 11% more distinct peptides, 1.2% more proteins and an increase from 49.5% to 51.6% in global proteome sequence coverage. Furthermore, this new PeptideAtlas release includes 5198 phosphorylated proteins, 668 ubiquitinated proteins, 3050 N-terminally acetylated proteins and 864 lysine-acetylated proteins. The majority of predicted Arabidopsis proteins has now been identified by MS, and users can explore the PeptideAtlas to readily determine if their proteins of interest have been identified, in which type of tissues or samples, obtain a sense of abundance, and evaluate if these proteins undergo any of the known major PTMs (phosphorylation, N-terminal or lysine acetylation, ubiquitination). Through GO enrichment analysis, machine learning, meta-data curation and analysis, as well as manual evaluation, we identified multiple reasons why proteins have not yet been identified in this new PeptideAtlas build. These reasons include i) small size (either because the gene encodes for a small protein or due to extensive proteolytic processing as in the case of signaling peptides), ii) high hydrophobicity, iii) very high pI, iv) low abundance (low expression or short-half-life), v) unusual PTMs, or vi) only presence in very specific conditions or cell types that were not included in the selected PXDs. The challenge now is to identify these remaining 20% of the predicted Arabidopsis proteome. Furthermore, this new build also mapped peptides to an additional ∼80 proteins not represented in the current Arabidopsis genome. These additional proteins should be considered in the community effort led by to Tanya Berardini at TAIR to generate a new annotation for Col-0 (tinyurl.com/Athalianav12).

This PeptideAtlas was built using about ∼20% of the currently (July 2022) available PXDs for Arabidopsis; incorporation of the vast majority of the unused PXDs is likely to only marginally increase the number of identified proteins as inferred from our comparison between build 1 and build 2. It is also not feasible to incorporate all these available raw data given the necessary time and expertise required. Furthermore, in case of several older PXDs in ProteomeXchange, low resolution instruments (*e.g*. LCQs or LTQs) or MALDI-TOF-TOF instruments were used; data from such PXDs are unlikely to contribute much to the PeptideAtlas (We note that even older data sets from 2005 – 2012 originally submitted to PRIDE are not available in ProteomeXchange).

To increase the number of protein identifications in PeptideAtlas, a strategic approach will be needed, by very carefully selecting data sets with the most sophisticated workflows (including selective enrichment for PTMs) and acquisition using the very latest generation of MS instruments (high mass accuracy, sensitivity and high dynamic range, very fast acquisition rates). Finally, a targeted approach to identify the missing (dark) proteome might be most effective using the combined insights from the machine learning models and the predicted protein properties and large-scale RNA-seq analysis across cell and tissue types, as well as developmental stages, biotic, and abiotic conditions.

## AUTHOR CONTRIBUTIONS

TL and ZS carried out the MS searches and PeptideAtlas data loading, supervised by EWD, and assembled the search results. ML developed the machine learning code. AK and AN assembled and analyzed the RNA-seq data. ML and SM contributed to data analysis and created figures. LM and ZS developed the PeptideAtlas web interface. IG, ED, GS, PR helped annotated the metadata in PeptideAtlas. QS helped assemble the protein search space. EWD and KJVW developed, coordinated and oversaw the project, evaluated outcomes, and wrote the paper.

## ACKNOWLEGDEMENTS

We thank members of the Scientific Advisory board of the NSF grant IOS-1922871 Tanya Berardini, Chris Town, Nicholas Provart, Sixue Chen and Joshua Heazlewood for advice and feedback.

## SUPPLEMENTAL DATA

**Supplemental Data Set S1.** Comprehensive overview of the 115 PXDs and their 369 experiments used for build 2. This includes key metadata as well as summaries of search results.

**Supplemental Data Set S2.** Transcript per million (TPM) expression values of 26975 predicted nuclear protein coding genes in Araport11 and the number of RNA-seq data sets (total 5673 filtered datasets) in which they are transcribed (A). Note that 398 genes were not transcribed (or available due to overlapping genes or sequence similarity (B) in any of the RNA-seq datasets and an additional 345 protein coding genes were never expressed above the median (C).

**Supplemental Data Set S3.** Proteins identified in non-Araport11 sources by hierarchy of sources (for hierarchy see Table 5)

**Supplemental Data Set S4.** Identification of N-terminal acetylation (NTA) sites in canonical proteins in PeptideAtlas. NTA sites per protein identifier. For each NTA site, the # of PSMs are listed at different PTM score interval (0.95 <p<0.99; 0.99 <p<1.0; no choice), as well as the sum of PSMs.

**Supplemental Data Set S5.** Identification of lysine acetylation (Kac) sites in canonical proteins in PeptideAtlas. A, For each Kac site, the # of PSMs are listed at different PTM score interval (0.95 <p<0.99; 0.99 <p<1.0; no choice), as well as the sum of PSMs. B, Non-redundant set of proteins with their number of observed Kac sites and total PSMs.

**Supplemental Data Set S6.** Identification of phosphorylation (S,T,Y) sites in canonical proteins in PeptideAtlas. A, Summarizing information of detected phosphorylation sites. For each p-site, the # of PSMs are listed at different PTM score interval (0.95 <p<0.99; 0.99 <p<1.0; no choice), as well as the sum of PSMs. B, Summarizing information of phosphorylated proteins with one or more phospho-sites.

**Supplemental Data Set S7.** Identification of Ubiquitination sites in canonical proteins in PeptideAtlas. A, Summarizing information of detected UBI sites. For each UBI-site, the # of PSMs are listed at different PTM score interval (0.95 <p<0.99; 0.99 <p<1.0; no choice), as well as the sum of PSMs. B, Summarizing information of detected UBI sites identified by both the GLY and diGLY method. For each UBI-site, the # of PSMs are listed at different PTM score interval (0.95 <p<0.99; 0.99 <p<1.0; no choice), as well as the sum of PSMs. C, Summarizing information of UBI proteins with one or more UBI sites.

**Supplemental Data Set S8.** Combined PTM results for the canonical proteins in PeptideAtlas with identified PTM sites for N-terminal acetylation, lysine acetylation, phosphorylation and/or ubiquitination. Listed are the protein identifiers and their annotations, NTA sites, K-ac sites, phosphor sites, UBI sites. Indicated are the amino acid residues position(s) for each PTM and total number of PSMs for each PTM across these positions.

**Supplemental Data Set S9.** Nuclear-encoded proteins Araport11 identifiers (26977) with annotations, protein properties, RNA-seq-based transcript information, machine learning predicted probability to be canonical and identification status in PeptideAtlas.

**Supplemental Data Set S10.** GO enrichment results of dark proteins. A,B. GO enrichments and associate genes identifiers for the dark proteins compared to all Arabidopsis proteins. Top20 most significant for BP and MF are listed. C,D. GO enrichment and associated gene identifies for the 222 outlier dark proteins compared to all 5595 dark proteins.

**Supplemental Data Set S11. P**roteins coding for signaling peptides, their annotations, physicochemical properties, RNA expression patterns and identification status in PeptideAtlas.

**Supplemental Figure S1**. Plotted values from Table 2. (A) For each of the 115 datasets in the build, there is no apparent correlation between the number between the identification efficiency (%spectra IDed) and the size of the experiment (spectra searched). (B) Displays a strong positive correlation, signifying the more spectra searched, the greater MS runs there are. (C) Shows a tight positive correlation, displaying the more spectra searched the higher the number of distinct peptides.

**Supplemental Figure S2**. Plotted values from Table 2. (A) For each 115 datasets in the build, there is no apparent correlation for the number of %spectra IDed vs spectra searched, between the identification efficiency and the size of the experiment. (B) Displays a strong positive correlation, signifying the more spectra searched, the greater MS runs there are. (C) Shows a tight positive correlation, displaying the more spectra searched the higher the number of distinct peptides. (D) A positive correlation between spectra searched and distinct canonical proteins can be observed.

## REFERENCES

Abbas M, Sharma G, Dambire C, Marquez J, Alonso-Blanco C, Proano K, Holdsworth MJ (2022) An oxygen-sensing mechanism for angiosperm adaptation to altitude. Nature 606: 565–569

Alex Mason G, Canto-Pastor A, Brady SM, Provart NJ (2021) Bioinformatic Tools in Arabidopsis Research. Methods Mol Biol 2200: 25–89

Ashburner M, Ball CA, Blake JA, Botstein D, Butler H, Cherry JM, Davis AP, Dolinski K, Dwight SS, Eppig JT, Harris MA, Hill DP, Issel-Tarver L, Kasarskis A, Lewis S, Matese JC, Richardson JE, Ringwald M, Rubin GM, Sherlock G (2000) Gene ontology: tool for the unification of biology. The Gene Ontology Consortium. Nat Genet 25: 25–29

Balparda M, Elsasser M, Badia MB, Giese J, Bovdilova A, Hudig M, Reinmuth L, Eirich J, Schwarzlander M, Finkemeier I, Schallenberg-Rudinger M, Maurino VG (2022) Acetylation of conserved lysines fine-tunes mitochondrial malate dehydrogenase activity in land plants. Plant J 109: 92–111

Barreto P, Dambire C, Sharma G, Vicente J, Osborne R, Yassitepe J, Gibbs DJ, Maia IG, Holdsworth MJ, Arruda P (2022) Mitochondrial retrograde signaling through UCP1- mediated inhibition of the plant oxygen-sensing pathway. Curr Biol 32: 1403–1411 e1404

Bartels S, Lori M, Mbengue M, van Verk M, Klauser D, Hander T, Boni R, Robatzek S, Boller T (2013) The family of Peps and their precursors in Arabidopsis: differential expression and localization but similar induction of pattern-triggered immune responses. J Exp Bot 64: 5309–5321

Bassal M, Abukhalaf M, Majovsky P, Thieme D, Herr T, Ayash M, Tabassum N, Al Shweiki MR, Proksch C, Hmedat A, Ziegler J, Lee J, Neumann S, Hoehenwarter W (2020) Reshaping of the Arabidopsis thaliana Proteome Landscape and Co-regulation of Proteins in Development and Immunity. Mol Plant 13: 1709–1732

Berger N, Vignols F, Przybyla-Toscano J, Roland M, Rofidal V, Touraine B, Zienkiewicz K, Couturier J, Feussner I, Santoni V, Rouhier N, Gaymard F, Dubos C (2020) Identification of client iron-sulfur proteins of the chloroplastic NFU2 transfer protein in Arabidopsis thaliana. J Exp Bot 71: 4171–4187

Bienvenut WV, Brunje A, Boyer JB, Muhlenbeck JS, Bernal G, Lassowskat I, Dian C, Linster E, Dinh TV, Koskela MM, Jung V, Seidel J, Schyrba LK, Ivanauskaite A, Eirich J, Hell R, Schwarzer D, Mulo P, Wirtz M, Meinnel T, Giglione C, Finkemeier I (2020) Dual lysine and N-terminal acetyltransferases reveal the complexity underpinning protein acetylation. Mol Syst Biol 16: e9464

Birnbaum KD, Otegui MS, Bailey-Serres J, Rhee SY (2022) The Plant Cell Atlas: focusing new technologies on the kingdom that nourishes the planet. Plant Physiol 188: 675–679

Chambers MC, Maclean B, Burke R, Amodei D, Ruderman DL, Neumann S, Gatto L, Fischer B, Pratt B, Egertson J, Hoff K, Kessner D, Tasman N, Shulman N, Frewen B, Baker TA, Brusniak MY, Paulse C, Creasy D, Flashner L, Kani K, Moulding C, Seymour SL, Nuwaysir LM, Lefebvre B, Kuhlmann F, Roark J, Rainer P, Detlev S, Hemenway T, Huhmer A, Langridge J, Connolly B, Chadick T, Holly K, Eckels J, Deutsch EW, Moritz RL, Katz JE, Agus DB, MacCoss M, Tabb DL, Mallick P (2012) A cross-platform toolkit for mass spectrometry and proteomics. Nat Biotechnol 30: 918–920

Chen X, Sun Y, Zhang T, Shu L, Roepstorff P, Yang F (2021) Quantitative Proteomics Using Isobaric Labeling: A Practical Guide. Genomics Proteomics Bioinformatics 19: 689–706

Cheng CY, Krishnakumar V, Chan AP, Thibaud-Nissen F, Schobel S, Town CD (2017) Araport11: a complete reannotation of the Arabidopsis thaliana reference genome. Plant J 89: 789–804

Chi W, He B, Mao J, Jiang J, Zhang L (2015) Plastid sigma factors: Their individual functions and regulation in transcription. Biochim Biophys Acta 1847: 770–778

Dahhan DA, Reynolds GD, Cardenas JJ, Eeckhout D, Johnson A, Yperman K, Kaufmann WA, Vang N, Yan X, Hwang I, Heese A, De Jaeger G, Friml J, Van Damme D, Pan J, Bednarek SY (2022) Proteomic characterization of isolated Arabidopsis clathrin-coated vesicles reveals evolutionarily conserved and plant-specific components. Plant Cell 34: 2150–2173

Deutsch EW, Bandeira N, Perez-Riverol Y, Sharma V, Carver JJ, Mendoza L, Kundu DJ, Wang S, Bandla C, Kamatchinathan S, Hewapathirana S, Pullman BS, Wertz J, Sun Z, Kawano S, Okuda S, Watanabe Y, MacLean B, MacCoss MJ, Zhu Y, Ishihama Y, Vizcaino JA (2023) The ProteomeXchange consortium at 10 years: 2023 update. Nucleic Acids Res 51: D1539–D1548

Deutsch EW, Mendoza L, Shteynberg D, Slagel J, Sun Z, Moritz RL (2015) Trans-Proteomic Pipeline, a standardized data processing pipeline for large-scale reproducible proteomics informatics. Proteomics Clin Appl 9: 745–754

Deutsch EW, Mendoza L, Shteynberg DD, Hoopmann MR, Sun Z, Eng JK, Moritz RL (2023) Trans-Proteomic Pipeline: Robust Mass Spectrometry-Based Proteomics Data Analysis Suite. J Proteome Res doi: 10.1021/acs.jproteome.2c00748. Online ahead of print.

Deutsch EW, Overall CM, Van Eyk JE, Baker MS, Paik YK, Weintraub ST, Lane L, Martens L, Vandenbrouck Y, Kusebauch U, Hancock WS, Hermjakob H, Aebersold R, Moritz RL, Omenn GS (2016) Human Proteome Project Mass Spectrometry Data Interpretation Guidelines 2.1. J Proteome Res 15: 3961–3970

Dinh TV, Bienvenut WV, Linster E, Feldman-Salit A, Jung VA, Meinnel T, Hell R, Giglione C, Wirtz M (2015) Molecular identification and functional characterization of the first Nalpha-acetyltransferase in plastids by global acetylome profiling. Proteomics 15: 2426–2435

Eng JK, Deutsch EW (2020) Extending Comet for Global Amino Acid Variant and Post-Translational Modification Analysis Using the PSI Extended FASTA Format. Proteomics 20: e1900362

Frankenfield AM, Ni J, Ahmed M, Hao L (2022) Protein Contaminants Matter: Building Universal Protein Contaminant Libraries for DDA and DIA Proteomics. J Proteome Res 21: 2104–2113

Fujita S (2021) CASPARIAN STRIP INTEGRITY FACTOR (CIF) family peptides - regulator of plant extracellular barriers. Peptides 143: 170599

Fussl M, Konig AC, Eirich J, Hartl M, Kleinknecht L, Bohne AV, Harzen A, Kramer K, Leister D, Nickelsen J, Finkemeier I (2022) Dynamic light- and acetate-dependent regulation of the proteome and lysine acetylome of Chlamydomonas. Plant J 109: 261–277

Ge SX, Jung D, Yao R (2020) ShinyGO: a graphical gene-set enrichment tool for animals and plants. Bioinformatics 36: 2628–2629

Gene Ontology C (2021) The Gene Ontology resource: enriching a GOld mine. Nucleic Acids Res 49: D325–D334

Gevaert K, Goethals M, Martens L, Van Damme J, Staes A, Thomas GR, Vandekerckhove J (2003) Exploring proteomes and analyzing protein processing by mass spectrometric identification of sorted N-terminal peptides. Nat Biotechnol 21: 566–569

Gibbs DJ, Conde JV, Berckhan S, Prasad G, Mendiondo GM, Holdsworth MJ (2015) Group VII Ethylene Response Factors Coordinate Oxygen and Nitric Oxide Signal Transduction and Stress Responses in Plants. Plant Physiol 169: 23–31

Giglione C, Boularot A, Meinnel T (2004) Protein N-terminal methionine excision. Cell Mol Life Sci 61: 1455–1474

Grubb LE, Derbyshire P, Dunning KE, Zipfel C, Menke FLH, Monaghan J (2021) Large-scale identification of ubiquitination sites on membrane-associated proteins in Arabidopsis thaliana seedlings. Plant Physiol 185: 1483–1488

Gunaratne J, Schmidt A, Quandt A, Neo SP, Sarac OS, Gracia T, Loguercio S, Ahrne E, Xia RL, Tan KH, Lossner C, Bahler J, Beyer A, Blackstock W, Aebersold R (2013) Extensive mass spectrometry-based analysis of the fission yeast proteome: the Schizosaccharomyces pombe PeptideAtlas. Mol Cell Proteomics 12: 1741–1751

Guo Y, Xiong L, Ishitani M, Zhu JK (2002) An Arabidopsis mutation in translation elongation factor 2 causes superinduction of CBF/DREB1 transcription factor genes but blocks the induction of their downstream targets under low temperatures. Proc Natl Acad Sci U S A 99: 7786–7791

Hains PG, Robinson PJ (2017) The Impact of Commonly Used Alkylating Agents on Artifactual Peptide Modification. J Proteome Res 16: 3443–3447

Hammarlund EU, Flashman E, Mohlin S, Licausi F (2020) Oxygen-sensing mechanisms across eukaryotic kingdoms and their roles in complex multicellularity. Science 370

Hawkins CL, Davies MJ (2019) Detection, identification, and quantification of oxidative protein modifications. J Biol Chem 294: 19683–19708

Hazarika RR, De Coninck B, Yamamoto LR, Martin LR, Cammue BP, van Noort V (2017) ARA-PEPs: a repository of putative sORF-encoded peptides in Arabidopsis thaliana. BMC Bioinformatics 18: 37

Hesselager MO, Codrea MC, Sun Z, Deutsch EW, Bennike TB, Stensballe A, Bundgaard L, Moritz RL, Bendixen E (2016) The Pig PeptideAtlas: A resource for systems biology in animal production and biomedicine. Proteomics 16: 634–644

Hodge K, Have ST, Hutton L, Lamond AI (2013) Cleaning up the masses: exclusion lists to reduce contamination with HPLC-MS/MS. J Proteomics 88: 92–103

Hooper CM, Castleden IR, Tanz SK, Aryamanesh N, Millar AH (2017) SUBA4: the interactive data analysis centre for Arabidopsis subcellular protein locations. Nucleic Acids Res 45: D1064–D1074

Hsu JL, Huang SY, Chow NH, Chen SH (2003) Stable-isotope dimethyl labeling for quantitative proteomics. Anal Chem 75: 6843–6852

Hu XL, Lu H, Hassan MM, Zhang J, Yuan G, Abraham PE, Shrestha HK, Villalobos Solis MI, Chen JG, Tschaplinski TJ, Doktycz MJ, Tuskan GA, Cheng ZM, Yang X (2021) Advances and perspectives in discovery and functional analysis of small secreted proteins in plants. Hortic Res 8: 130

Huang A, Tang Y, Shi X, Jia M, Zhu J, Yan X, Chen H, Gu Y (2020) Proximity labeling proteomics reveals critical regulators for inner nuclear membrane protein degradation in plants. Nat Commun 11: 3284

Huang S, Taylor NL, Whelan J, Millar AH (2009) Refining the definition of plant mitochondrial presequences through analysis of sorting signals, N-terminal modifications, and cleavage motifs. Plant Physiol 150: 1272–1285

Huffaker A, Pearce G, Ryan CA (2006) An endogenous peptide signal in Arabidopsis activates components of the innate immune response. Proc Natl Acad Sci U S A 103: 10098–10103

Hulstaert N, Shofstahl J, Sachsenberg T, Walzer M, Barsnes H, Martens L, Perez-Riverol Y (2020) ThermoRawFileParser: Modular, Scalable, and Cross-Platform RAW File Conversion. J Proteome Res 19: 537–542

Kaufmann C, Sauter M (2019) Sulfated plant peptide hormones. J Exp Bot 70: 4267–4277

Keller A, Eng J, Zhang N, Li XJ, Aebersold R (2005) A uniform proteomics MS/MS analysis platform utilizing open XML file formats. Mol Syst Biol 1: 2005 0017

Keller A, Nesvizhskii AI, Kolker E, Aebersold R (2002) Empirical statistical model to estimate the accuracy of peptide identifications made by MS/MS and database search. Anal Chem 74: 5383–5392

Kim JS, Jeon BW, Kim J (2021) Signaling Peptides Regulating Abiotic Stress Responses in Plants. Front Plant Sci 12: 704490

Kim MS, Zhong J, Pandey A (2016) Common errors in mass spectrometry-based analysis of post-translational modifications. Proteomics 16: 700–714

King NL, Deutsch EW, Ranish JA, Nesvizhskii AI, Eddes JS, Mallick P, Eng J, Desiere F, Flory M, Martin DB, Kim B, Lee H, Raught B, Aebersold R (2006) Analysis of the Saccharomyces cerevisiae proteome with PeptideAtlas. Genome Biol 7: R106

Kleifeld O, Doucet A, Prudova A, auf dem Keller U, Gioia M, Kizhakkedathu JN, Overall CM (2011) Identifying and quantifying proteolytic events and the natural N terminome by terminal amine isotopic labeling of substrates. Nat Protoc 6: 1578-1611

Kong AT, Leprevost FV, Avtonomov DM, Mellacheruvu D, Nesvizhskii AI (2017) MSFragger: ultrafast and comprehensive peptide identification in mass spectrometry-based proteomics. Nat Methods 14: 513–520

Koornneef M, Meinke D (2011) The development of Arabidopsis as a model plant. Plant J 61: 909–921

Kyte J, Doolittle RF (1982) A simple method for displaying the hydropathic character of a protein. J Mol Biol 157: 105–132

Lamesch P, Berardini TZ, Li D, Swarbreck D, Wilks C, Sasidharan R, Muller R, Dreher K, Alexander DL, Garcia-Hernandez M, Karthikeyan AS, Lee CH, Nelson WD, Ploetz L, Singh S, Wensel A, Huala E (2012) The Arabidopsis Information Resource (TAIR): improved gene annotation and new tools. Nucleic Acids Res 40: D1202–1210

Li W, O’Neill KR, Haft DH, DiCuccio M, Chetvernin V, Badretdin A, Coulouris G, Chitsaz F, Derbyshire MK, Durkin AS, Gonzales NR, Gwadz M, Lanczycki CJ, Song JS, Thanki N, Wang J, Yamashita RA, Yang M, Zheng C, Marchler-Bauer A, Thibaud-Nissen F (2021) RefSeq: expanding the Prokaryotic Genome Annotation Pipeline reach with protein family model curation. Nucleic Acids Res 49: 1020–1028

Liao Y, Smyth GK, Shi W (2014) featureCounts: an efficient general purpose program for assigning sequence reads to genomic features. Bioinformatics 30: 923–930

Ma J, Chen T, Wu S, Yang C, Bai M, Shu K, Li K, Zhang G, Jin Z, He F, Hermjakob H, Zhu Y (2019) iProX: an integrated proteome resource. Nucleic Acids Res 47: D1211–D1217

Maddelein D, Colaert N, Buchanan I, Hulstaert N, Gevaert K, Martens L (2015) The iceLogo web server and SOAP service for determining protein consensus sequences. Nucleic Acids Res 43: W543–546

Malmstrom J, Beck M, Schmidt A, Lange V, Deutsch EW, Aebersold R (2009) Proteome-wide cellular protein concentrations of the human pathogen Leptospira interrogans. Nature 460: 762–765

Martens L, Chambers M, Sturm M, Kessner D, Levander F, Shofstahl J, Tang WH, Rompp A, Neumann S, Pizarro AD, Montecchi-Palazzi L, Tasman N, Coleman M, Reisinger F, Souda P, Hermjakob H, Binz PA, Deutsch EW (2011) mzML--a community standard for mass spectrometry data. Mol Cell Proteomics 10: R110 000133

Matsubayashi Y (2014) Posttranslationally modified small-peptide signals in plants. Annu Rev Plant Biol 65: 385–413

McCord J, Sun Z, Deutsch EW, Moritz RL, Muddiman DC (2017) The PeptideAtlas of the Domestic Laying Hen. J Proteome Res 16: 1352–1363

Medina J, Ballesteros ML, Salinas J (2007) Phylogenetic and functional analysis of Arabidopsis RCI2 genes. J Exp Bot 58: 4333–4346

Meinke DW, Cherry JM, Dean C, Rounsley SD, Koornneef M (1998) Arabidopsis thaliana: a model plant for genome analysis. Science 282: 662, 679–682

Meinnel T, Giglione C (2022) N-terminal modifications, the associated processing machinery, and their evolution in plastid-containing organisms. J Exp Bot 73: 6013–6033

Mergner J, Frejno M, List M, Papacek M, Chen X, Chaudhary A, Samaras P, Richter S, Shikata H, Messerer M, Lang D, Altmann S, Cyprys P, Zolg DP, Mathieson T, Bantscheff M, Hazarika RR, Schmidt T, Dawid C, Dunkel A, Hofmann T, Sprunck S, Falter-Braun P, Johannes F, Mayer KFX, Jurgens G, Wilhelm M, Baumbach J, Grill E, Schneitz K, Schwechheimer C, Kuster B (2020) Mass-spectrometry-based draft of the Arabidopsis proteome. Nature 579: 409–414

Michalik S, Depke M, Murr A, Gesell Salazar M, Kusebauch U, Sun Z, Meyer TC, Surmann K, Pfortner H, Hildebrandt P, Weiss S, Palma Medina LM, Gutjahr M, Hammer E, Becher D, Pribyl T, Hammerschmidt S, Deutsch EW, Bader SL, Hecker M, Moritz RL, Mader U, Volker U, Schmidt F (2017) A global Staphylococcus aureus proteome resource applied to the in vivo characterization of host-pathogen interactions. Sci Rep 7: 9718

Moriya Y, Kawano S, Okuda S, Watanabe Y, Matsumoto M, Takami T, Kobayashi D, Yamanouchi Y, Araki N, Yoshizawa AC, Tabata T, Iwasaki M, Sugiyama N, Tanaka S, Goto S, Ishihama Y (2019) The jPOST environment: an integrated proteomics data repository and database. Nucleic Acids Res 47: D1218–D1224

Muller T, Winter D (2017) Systematic Evaluation of Protein Reduction and Alkylation Reveals Massive Unspecific Side Effects by Iodine-containing Reagents. Mol Cell Proteomics 16: 1173–1187

Nissa MU, Reddy PJ, Pinto N, Sun Z, Ghosh B, Moritz RL, Goswami M, Srivastava S (2022) The PeptideAtlas of a widely cultivated fish Labeo rohita: A resource for the Aquaculture Community. Sci Data 9: 171

Niu B, Martinelli Ii M, Jiao Y, Wang C, Cao M, Wang J, Meinke E (2020) Nonspecific cleavages arising from reconstitution of trypsin under mildly acidic conditions. PLoS One 15: e0236740

Olsson V, Joos L, Zhu S, Gevaert K, Butenko MA, De Smet I (2019) Look Closely, the Beautiful May Be Small: Precursor-Derived Peptides in Plants. Annu Rev Plant Biol 70: 153–186

Omenn GS, Lane L, Overall CM, Paik YK, Cristea IM, Corrales FJ, Lindskog C, Weintraub S, Roehrl MHA, Liu S, Bandeira N, Srivastava S, Chen YJ, Aebersold R, Moritz RL, Deutsch EW (2021) Progress Identifying and Analyzing the Human Proteome: 2021 Metrics from the HUPO Human Proteome Project. J Proteome Res 20: 5227–5240

Palos K, Nelson Dittrich AC, Yu L, Brock JR, Railey CE, Wu HL, Sokolowska E, Skirycz A, Hsu PY, Gregory BD, Lyons E, Beilstein MA, Nelson ADL (2022) Identification and functional annotation of long intergenic non-coding RNAs in Brassicaceae. Plant Cell 34: 3233–3260

Parry G, Provart NJ, Brady SM, Uzilday B, Multinational Arabidopsis Steering C (2020) Current status of the multinational Arabidopsis community. Plant Direct 4: e00248

Perez-Riverol Y, Bai J, Bandla C, Garcia-Seisdedos D, Hewapathirana S, Kamatchinathan S, Kundu DJ, Prakash A, Frericks-Zipper A, Eisenacher M, Walzer M, Wang S, Brazma A, Vizcaino JA (2022) The PRIDE database resources in 2022: a hub for mass spectrometry-based proteomics evidences. Nucleic Acids Res 50: D543–D552

Perez-Riverol Y, Csordas A, Bai J, Bernal-Llinares M, Hewapathirana S, Kundu DJ, Inuganti A, Griss J, Mayer G, Eisenacher M, Perez E, Uszkoreit J, Pfeuffer J, Sachsenberg T, Yilmaz S, Tiwary S, Cox J, Audain E, Walzer M, Jarnuczak AF, Ternent T, Brazma A, Vizcaino JA (2018) The PRIDE database and related tools and resources in 2019: improving support for quantification data. Nucleic Acids Res 47: D442–D450

Plant Cell Atlas C, Jha SG, Borowsky AT, Cole BJ, Fahlgren N, Farmer A, Huang SC, Karia P, Libault M, Provart NJ, Rice SL, Saura-Sanchez M, Agarwal P, Ahkami AH, Anderton CR, Briggs SP, Brophy JA, Denolf P, Di Costanzo LF, Exposito-Alonso M, Giacomello S, Gomez-Cano F, Kaufmann K, Ko DK, Kumar S, Malkovskiy AV, Nakayama N, Obata T, Otegui MS, Palfalvi G, Quezada-Rodriguez EH, Singh R, Uhrig RG, Waese J, Van Wijk K, Wright RC, Ehrhardt DW, Birnbaum KD, Rhee SY (2021) Vision, challenges and opportunities for a Plant Cell Atlas. Elife 10

Pozoga M, Armbruster L, Wirtz M (2022) From Nucleus to Membrane: A Subcellular Map of the N-Acetylation Machinery in Plants. Int J Mol Sci 23

Provart NJ, Brady SM, Parry G, Schmitz RJ, Queitsch C, Bonetta D, Waese J, Schneeberger K, Loraine AE (2021) Anno genominis XX: 20 years of Arabidopsis genomics. Plant Cell 33: 832–845

Pullman BS, Wertz J, Carver J, Bandeira N (2018) ProteinExplorer: A Repository-Scale Resource for Exploration of Protein Detection in Public Mass Spectrometry Data Sets. J Proteome Res 17: 4227–4234

Puthiyaveetil S, McKenzie SD, Kayanja GE, Ibrahim IM (2021) Transcription initiation as a control point in plastid gene expression. Biochim Biophys Acta Gene Regul Mech 1864: 194689

Rauniyar N, Yates JR, 3rd (2014) Isobaric labeling-based relative quantification in shotgun proteomics. J Proteome Res 13: 5293–5309

Reales-Calderon JA, Sun Z, Mascaraque V, Perez-Navarro E, Vialas V, Deutsch EW, Moritz RL, Gil C, Martinez JL, Molero G (2021) A wide-ranging Pseudomonas aeruginosa PeptideAtlas build: A useful proteomic resource for a versatile pathogen. J Proteomics 239: 104192

Rodriguez E, Chevalier J, Olsen J, Ansbol J, Kapousidou V, Zuo Z, Svenning S, Loefke C, Koemeda S, Drozdowskyj PS, Jez J, Durnberger G, Kuenzl F, Schutzbier M, Mechtler K, Ebstrup EN, Lolle S, Dagdas Y, Petersen M (2020) Autophagy mediates temporary reprogramming and dedifferentiation in plant somatic cells. EMBO J 39: e103315

Ross S, Giglione C, Pierre M, Espagne C, Meinnel T (2005) Functional and developmental impact of cytosolic protein N-terminal methionine excision in Arabidopsis. Plant Physiol 137: 623–637

Rowland E, Kim J, Bhuiyan NH, van Wijk KJ (2015) The Arabidopsis Chloroplast Stromal N-Terminome: Complexities of Amino-Terminal Protein Maturation and Stability. Plant Physiol 169: 1881–1896

Rytz TC, Miller MJ, McLoughlin F, Augustine RC, Marshall RS, Juan YT, Charng YY, Scalf M, Smith LM, Vierstra RD (2018) SUMOylome Profiling Reveals a Diverse Array of Nuclear Targets Modified by the SUMO Ligase SIZ1 during Heat Stress. Plant Cell 30: 1077–1099

Sanderfoot AA, Kovaleva V, Zheng H, Raikhel NV (1999) The t-SNARE AtVAM3p resides on the prevacuolar compartment in Arabidopsis root cells. Plant Physiol 121: 929–938

Schittmayer M, Fritz K, Liesinger L, Griss J, Birner-Gruenberger R (2016) Cleaning out the Litterbox of Proteomic Scientists’ Favorite Pet: Optimized Data Analysis Avoiding Trypsin Artifacts. J Proteome Res 15: 1222–1229

Sharma V, Eckels J, Schilling B, Ludwig C, Jaffe JD, MacCoss MJ, MacLean B (2018) Panorama Public: A Public Repository for Quantitative Data Sets Processed in Skyline. Mol Cell Proteomics 17: 1239–1244

Shteynberg D, Deutsch EW, Lam H, Eng JK, Sun Z, Tasman N, Mendoza L, Moritz RL, Aebersold R, Nesvizhskii AI (2011) iProphet: multi-level integrative analysis of shotgun proteomic data improves peptide and protein identification rates and error estimates. Mol Cell Proteomics 10: M111 007690

Shteynberg DD, Deutsch EW, Campbell DS, Hoopmann MR, Kusebauch U, Lee D, Mendoza L, Midha MK, Sun Z, Whetton AD, Moritz RL (2019) PTMProphet: Fast and Accurate Mass Modification Localization for the Trans-Proteomic Pipeline. J Proteome Res 18: 4262–4272

Silva J, Ferraz R, Dupree P, Showalter AM, Coimbra S (2020) Three Decades of Advances in Arabinogalactan-Protein Biosynthesis. Front Plant Sci 11: 610377

Sloan DB, Wu Z, Sharbrough J (2018) Correction of Persistent Errors in Arabidopsis Reference Mitochondrial Genomes. Plant Cell 30: 525–527

Somerville CR, Ogren WL (1980) Inhibition of photosynthesis in Arabidopsis mutants lacking leaf glutamate synthase activity. Nature 286: 257–259

Somerville CR, Ogren WL (1982) Mutants of the cruciferous plant Arabidopsis thaliana lacking glycine decarboxylase activity. Biochem J 202: 373–380

Stintzi A, Schaller A (2022) Biogenesis of post-translationally modified peptide signals for plant reproductive development. Curr Opin Plant Biol 69: 102274

Sun Q, Zybailov B, Majeran W, Friso G, Olinares PD, van Wijk KJ (2009) PPDB, the Plant Proteomics Database at Cornell. Nucleic Acids Res 37: D969–974

Takahashi F, Hanada K, Kondo T, Shinozaki K (2019) Hormone-like peptides and small coding genes in plant stress signaling and development. Curr Opin Plant Biol 51: 88–95

Tavormina P, De Coninck B, Nikonorova N, De Smet I, Cammue BP (2015) The Plant Peptidome: An Expanding Repertoire of Structural Features and Biological Functions. Plant Cell 27: 2095–2118

Tilak P, Kotnik F, Nee G, Seidel J, Sindlinger J, Heinkow P, Eirich J, Schwarzer D, Finkemeier I (2023) Proteome-wide lysine acetylation profiling to investigate the involvement of histone deacetylase HDA5 in the salt stress response of Arabidopsis leaves. Plant J

Tost AS, Kristensen A, Olsen LI, Axelsen KB, Fuglsang AT (2021) The PSY Peptide Family-Expression, Modification and Physiological Implications. Genes (Basel) 12

UniProt C (2020) UniProt: the universal protein knowledgebase in 2021. Nucleic Acids Res

UniProt C (2023) UniProt: the Universal Protein Knowledgebase in 2023. Nucleic Acids Res 51: D523–D531

van Dongen JT, Licausi F (2015) Oxygen sensing and signaling. Annu Rev Plant Biol 66: 345–367

van Wijk KJ, Friso G, Walther D, Schulze WX (2014) Meta-Analysis of Arabidopsis thaliana Phospho-Proteomics Data Reveals Compartmentalization of Phosphorylation Motifs. Plant Cell 26: 2367–2389

van Wijk KJ, Leppert T, Sun Q, Boguraev SS, Sun Z, Mendoza L, Deutsch EW (2021) The Arabidopsis PeptideAtlas: Harnessing worldwide proteomics data to create a comprehensive community proteomics resource. Plant Cell 33: 3421–3453

Verrastro I, Pasha S, Jensen KT, Pitt AR, Spickett CM (2015) Mass spectrometry-based methods for identifying oxidized proteins in disease: advances and challenges. Biomolecules 5: 378–411

Walton A, Stes E, Cybulski N, Van Bel M, Inigo S, Durand AN, Timmerman E, Heyman J, Pauwels L, De Veylder L, Goossens A, De Smet I, Coppens F, Goormachtig S, Gevaert K (2016) It’s Time for Some “Site”-Seeing: Novel Tools to Monitor the Ubiquitin Landscape in Arabidopsis thaliana. Plant Cell 28: 6–16

Waltz F, Nguyen TT, Arrive M, Bochler A, Chicher J, Hammann P, Kuhn L, Quadrado M, Mireau H, Hashem Y, Giege P (2019) Small is big in Arabidopsis mitochondrial ribosome. Nat Plants 5: 106–117

Weits DA, van Dongen JT, Licausi F (2021) Molecular oxygen as a signaling component in plant development. New Phytol 229: 24–35

White MD, Klecker M, Hopkinson RJ, Weits DA, Mueller C, Naumann C, O’Neill R, Wickens J, Yang J, Brooks-Bartlett JC, Garman EF, Grossmann TN, Dissmeyer N, Flashman E (2017) Plant cysteine oxidases are dioxygenases that directly enable arginyl transferase-catalysed arginylation of N-end rule targets. Nat Commun 8: 14690

Willems P (2022) Exploring Posttranslational Modifications with the Plant PTM Viewer. Methods Mol Biol 2447: 285–296

Willems P, Ndah E, Jonckheere V, Van Breusegem F, Van Damme P (2021) To New Beginnings: Riboproteogenomics Discovery of N-Terminal Proteoforms in Arabidopsis Thaliana. Front Plant Sci 12: 778804

Willoughby AC, Nimchuk ZL (2021) WOX going on: CLE peptides in plant development. Curr Opin Plant Biol 63: 102056

Wu GZ, Bock R (2021) GUN control in retrograde signaling: How GENOMES UNCOUPLED proteins adjust nuclear gene expression to plastid biogenesis. Plant Cell 33: 457–474

Yuan B, Wang H (2021) Peptide Signaling Pathways Regulate Plant Vascular Development. Front Plant Sci 12: 719606

Zhang M, Tan FQ, Fan YJ, Wang TT, Song X, Xie KD, Wu XM, Zhang F, Deng XX, Grosser JW, Guo WW (2022) Acetylome reprograming participates in the establishment of fruit metabolism during polyploidization in citrus. Plant Physiol 190: 2519–2538

Zhong S, Liu M, Wang Z, Huang Q, Hou S, Xu YC, Ge Z, Song Z, Huang J, Qiu X, Shi Y, Xiao J, Liu P, Guo YL, Dong J, Dresselhaus T, Gu H, Qu LJ (2019) Cysteine-rich peptides promote interspecific genetic isolation in Arabidopsis. Science 364

Zybailov B, Sun Q, van Wijk KJ (2009) Workflow for large scale detection and validation of peptide modifications by RPLC-LTQ-Orbitrap: application to the Arabidopsis thaliana leaf proteome and an online modified peptide library. Anal Chem 81: 8015–8024

